# Anatomically-specific coupling between innate immune gene repertoire and microbiome structure during coral evolution

**DOI:** 10.1101/2023.04.26.538298

**Authors:** Tanya Brown, Dylan Sonett, Ryan McMinds, F. Joseph Pollock, Mónica Medina, Jesse R. Zaneveld

## Abstract

Tropical reef-building corals exist in intimate symbiosis with diverse microbes and viruses. Coral microbiomes are generally much less diverse than their environment, but across studied corals, the biodiversity of these microbiomes varies greatly. It has previously been hypothesized that differences in coral innate immunity in general, and the copy number of TIR-domain containing innate immune genes in particular, may drive interspecific differences in microbiome structure. Despite many existing studies of coral microbiomes, this hypothesis has previously been difficult to test due to a lack of consistently collected cross-species data on coral microbiomes. In this manuscript, we reannotate TIR-domain containing genes across diverse coral genomes, and use phylogenetic comparative methods to compare these innate immune gene copy numbers against 16S rRNA marker gene data on coral mucus, tissue, and skeleton microbiomes from the Global Coral Microbiome Project (GCMP). The copy number of Toll-like receptor (TLRs) and Interleukin-1 receptor (IL-1Rs) gene families, as well as the total genomic count of their constituent domains (LRR and TIR domains; and Ig and TIR domains, respectively), explained most interspecific differences in microbiome richness and beta-diversity among corals with sequenced genomes. We find that these correlations are also anatomically specific, with an especially strong correlation between IL-1R gene copy numbers and microbiome richness in the coral’s endolithic skeleton. Together, these results suggest innate immunity may play a key role in sculpting microbiome structure in corals.

## Introduction

The 1681 described species of scleractinian corals^1^ are environmentally critical ecosystem engineers that underpin many tropical reef ecosystems. Microbiomes are important contributors to the health of these tropical corals, with competing and cooperating microbes influencing animal health^2^. Therefore, evaluating how corals regulate this microbiome is of great importance. Numerous studies have uncovered important features of coral microbiomes, including the relative influence of differences across host anatomy^3,4^, between species^4,5^, among reefs^6–8^, and along environmental gradients^8,9^. This literature has also extensively documented coral microbiome responses to various stressors, including heat^10,11^, bleaching^12,13^, sedimentation^14,15^, nutrient pollution^10,14,16^, predation^16^, plastic pollution^17^, turf or macro-algal competition^16,18,19^, etc. Genetic studies within coral species have further found genotypic differences that correlate with microbiome composition^20,21^. Building on these ecological and population-genetic comparisons, specific coral microorganisms have been linked to important host health outcomes, such as protection against pathogens or susceptibility to them^22^. Recent microbiome manipulation experiments have even begun to establish the causal role of specific coral-associated bacteria in influencing key host traits like heat resistance^23^. Despite this thriving literature on coral microbiomes, the broader scale patterns of how modern coral microbiomes have evolved, and which host traits, if any, drive the large differences in microbiome structure and function seen between modern corals is not yet clear.

Increased attention to the question of how host traits have sculpted coral microbiomes over evolution is important. Comparative studies of coral microbiome evolution may identify host traits that have regulated coral microbiomes up to the present day. Comparative studies of coral microbiome evolution will also clarify a key part of the broader story of animal microbiome evolution. While vertebrate gut microbiomes are structured by both host phylogenetic relatedness and convergently evolved host traits like diet or flight^24,25^, the traits shaping the microbiome evolution of basally divergent animal taxa are less clear.

Among the many coral traits that could influence microbiome structure, differences in innate immunity are promising candidates, since they have the potential to directly regulate the microbiome by activating pathways that preferentially target particular groups of microbes. The copy number of gene families containing Toll/Interleukin Repeat (TIR) domains in particular, have received special attention as a possible influence on microbiome structure^26^. TIR domains are a key intracellular signaling domain, found in multiple innate immune gene families, such as Toll-like Receptors (TLR), Interleukin-1 Receptors (IL-1R), coral-specific TIR-only genes of unknown function, and Myeloid Differentiation Factor 88 (myD88). Collectively, these genes are known as TIR-domain containing genes^27^. The genomic copy number of some gene families of TIR-domain containing genes is known to vary greatly between coral species, and on this basis these genes have been hypothesized to influence cross-species differences in coral microbiome structure^26^. However, a lack of consistently collected cross-species microbiome data has so far precluded empirical testing of this intriguing idea.

In this study we tested whether dramatic changes in the copy number of coral TLR or IL-1R gene families have driven corresponding changes in microbiome richness, evenness, or composition during more than 250 million years of scleractinian coral evolution. Our analysis combined genomic analysis of all major lineages of scleractinian coral for which genomes are publicly available with data on coral mucus, tissue, and endolithic skeleton microbiomes from the Global Coral Microbiome Project (GCMP) dataset^3^ — a collection of more than 1440 16S rRNA libraries from diverse coral species. We then used established phylogenetic comparative methods to test whether coral species’ TLR or IL-1R gene families correlated with their microbiome structure or composition. The results identify potential drivers of microbiome structure among sequenced corals, and highlight the value of comparative analysis for supporting or refuting whether specific host traits underlie differences in microbiome structure between animal species. They further support the idea that different regions of coral anatomy differ not just in microbiome composition, but also in responsiveness to host traits — with the parts of the coral that are most strongly linked to host traits sometimes being a surprise.

## Methods

### Genomic Data Acquisition

Reference coral genomes were selected based on completeness and overlap with the Global Coral Microbiome Project dataset. Twelve coral genomes were downloaded from NCBI (https://www.ncbi.nlm.nih.gov/) and Reef Genomics (http://reefgenomics.org). A list of urls for the genomes can be found in **Table S1A**.

### Domain Annotation

The downloaded coral genomes were used to locate TIR, leucine rich repeats (LRR), and immunoglobulin (Ig) containing genes using a custom pipeline found on github (https://github.com/zaneveld/GCMP_genomics). Genome analysis occurred in two steps. First, genomes were analyzed using TransDecoder (v5.5.0) (https://github.com/TransDecoder/TransDecoder/wiki) to identify candidate coding regions within the genome files. During this process, TransDecoder converts the nucleotide sequences in the genome file to possible amino acid sequences using different open reading frames needed to conduct this translation. The peptide file generated from TransDecoder was next searched with HMMER (hmmer.org, HMMER 3.3.2 (November 2020)) to locate putative TIR domains as well as known TIR-associated domains. The TIR, LRR, and Ig associated peptide alignment files used in this search were downloaded from the Pfam website (http://pfam.xfam.org/). These 14 alignment files used are found in **Table S2B, C**. The Pfam domain alignment files were used to build profiles in HMMER using hmmbuild by reading in the alignment file and creating a new Hidden Markov Model (HMM) profile. The HMM profiles were then used to search the TransDecoder peptide files. The output file from hmmscan contained significant matches between the HMM profile and the transdecoder peptide file. The resultant hmmscan file was used to count the number of TIR-domain containing sequences found for each organism.

TIR-domain containing genes were further subdivided by scanning each for LRR or Ig domains using the same procedure. TIR-domain-containing genes were then subdivided into categories based on their domains: TIR only (defined by TIR domain and no LRR or Ig domains), TLR (TIR and one or more LRR domains), and IL-1R (TLR and one or more Ig domains) genes. Numbers of each type of gene, and average numbers of Ig or LRR domains within each type of gene were used in further analysis.

### Coral sampling and microbiome analysis

The Global Coral Microbiome Project (GCMP) coral mucus, tissue and skeleton samples reanalyzed here were originally collected and processed following the methods outlined in Pollock *et al.,* 2018^3^. Those methods are briefly restated here. All coral samples were collected by AAUS-certified scientific divers, in accordance with local regulations. These plus additional international samples were then resequenced with protocols standard for the Earth Microbiome Project^28^. Bacterial and archaeal DNA were extracted using the PowerSoil DNA Isolation Kit (MoBio Laboratories, Carlsbad, CA; now Qiagen, Venlo, Netherlands). To select for the 16S rRNA V4 gene region, polymerase chain reaction (PCR) was performed using the following primers with illumina adapter sequences (underlined) at the 5’ ends: 515F^29^ 5′− <u>TCG TCG GCA GCG TCA GAT GTG TAT</u> <u>AAG AGA CAG</u> GTG YCA GCM GCC GCG GTA A −3′ and 806R^30^ 5’− <u>GTC TCG</u> <u>TGG GCT CGG AGA TGT GTA TAA GAG ACA G</u>GG ACT CAN VGG GTW TCT AAT −3′. PCR, library preparation and sequencing on an Illumina HiSeq (2×125bp) was performed by the EMP^28,31^. The resulting 16S rRNA amplicon sequences from the GCMP are available in the Qiita database Qiita (CRC32 id: 8817b8b8 and CRC32 id: ac925c85). Metadata for all GCMP samples are available in **Table S1B**.

### 16S library preparation, sequencing, and initial quality control

16S rRNA sequencing data were processed in Qiita^32^ using the standard EMP workflow. Briefly, sequences were demultiplexed based on 12bp Golay barcodes using “split_libraries” with QIIME 1.9.1 default parameters^33^ and trimmed to 100bp to remove low quality base pairs. Quality control (e.g., denoising, de-replication and chimera filtering) and identification amplicon sequence variants (ASVs) were performed using deblur 1.1.0^34^ with default parameters. The resulting biom and taxonomy tables were obtained from Qiita (CRC32 id: 8817b8b8 and CRC32 id: ac925c85) and processed using a customized QIIME2 v. 2020.8.0 ^35^ pipeline in python (github.com/zaneveld/GCMP_global_disease).

### Mitochondrial annotation and quality control

Taxonomic assignment of ASVs was performed using vsearch^36^ using a modified version of the SILVA v. 138^37^ taxonomic reference. The sole change to the SILVA138 reference was supplementation with additional coral mitochondrial reads obtained from metaxa2^38^. In benchmarks, this change greatly improves annotation of coral mitochondrial rRNAs, without increasing false positive taxonomic assignments^39^. This expanded taxonomy is referred to as “silva_metaxa2” in code. After taxonomic assignment, all mitochondrial and chloroplast reads were removed (**Table S3**).

The bacterial phylogenetic tree was built using the SATé-enabled phylogenetic placement (SEPP) insertion technique with the q2-fragment-insertion plugin^40^, again using the SILVA v. 138^37^ database as reference taxonomy. The final output from this pipeline consisted of a taxonomy table, ASV feature table and phylogenetic tree that were used for downstream analyses.

Phylogenetic comparison of innate immune gene repertoire and microbiome richness Genome TIR domains were compared to the microbiome data collected from the GCMP. The α and < -diversity of the microbial community were compared to the TIR, TLR, and IL-R domain copy numbers using phylogenetically independent contrasts (PICs) with the phytools package in R. PICs use the coral species phylogenetic information in comparison with the TIR-associated domains and ASV and Gini Index information to assess the relatedness for α-diversity. PIC analysis for phylogenetic < -diversity metrics (Weighted and Unweighted UniFrac) were created from three PC axes (PC1, PC2, and PC3) of a PCoA ordination. Analyses were conducted on all three axes. The PIC method takes into account a priori the importance of phylogenetic history on trait variation when extracting information in relation to the common ancestor. Code used for this analysis is found on github.

### Microbial taxonomic analysis

Microbial taxonomic analysis was conducted using the ANCOM-BC package in phyloseq in R^41^. Analyses were conducted at the class and family level with relation to IL-1R and TLR copy numbers. Lists of significant microbes were generated for mucus, tissue, skeleton, and all compartments. Heatmaps were generated to visualize positive and negative correlation of the microbial community.

## Results

Sequenced coral genomes vary greatly in TIR-domain containing gene copy number. Previous analyses have reported significant variation in the copy number of TIR-domain containing genes among coral genomes. We annotated the genomic copy number of TIR-domain containing genes in 11 coral genomes that were also represented in microbiome data from the Global Coral Microbiome Project (GCMP) (**Table S1**). While some prior studies have analyzed both genomes and transcriptomes in order to maximize discovery of new TLR or IL-1R homologs, we chose to exclude transcriptomes from our analysis in order to prevent any potential confounding effects of some innate immune genes not being expressed in transcriptomes. As a result of these annotations (Methods), we identified numerous TLR, IL-1R and TIR-only genes (**Table S2, 4**). Despite these methodological differences, our annotations mostly agree in trend and rank with prior studies of coral innate immune repertoires^26,42^.

Many innate immune genes have modular structures based on the domains that they contain, and TIR domains commonly co-occur with several other domains within innate immune genes. We further subdivided TIR-domain containing genes based on the other domains present: genes with both TIR and immunoglobulin (Ig) domains were annotated as interleukin-1 receptors (IL-1R) while genes with both TIR domains and leucine rich repeat (LRR) domains were annotated as toll-like receptors (TLR) (**Fig. 1**). We also analyzed the total count of TIR domains, regardless of which gene they were part of and the presence or absence of other co-associated domains.

**Fig. 1.**
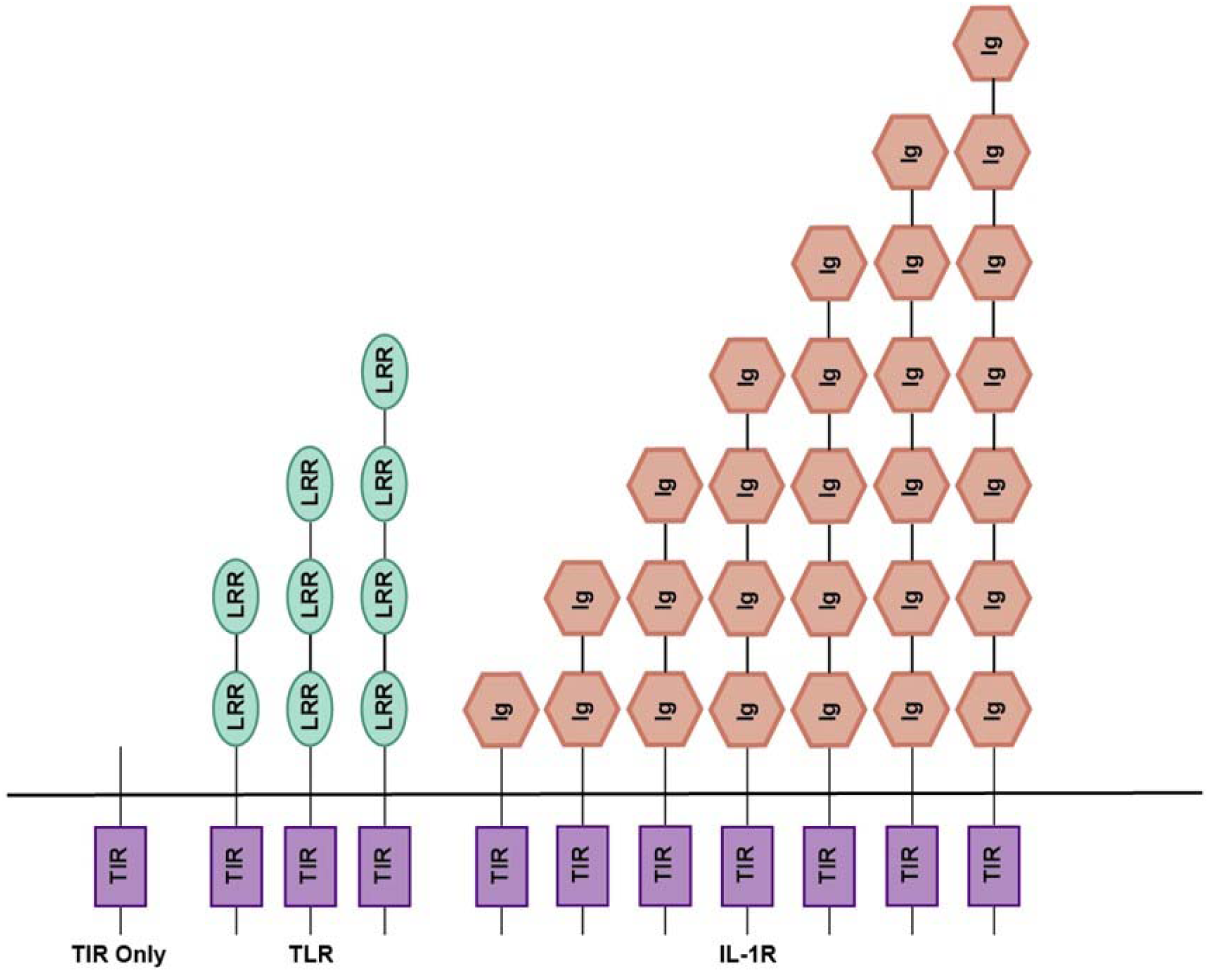
TIR containing gene architecture in coral genomes. Diagram depicts all domains present in TIR only, TLR, or IL-1R genes identified in this study. TIR: Toll-Interleukin Repeat; Ig: Immunoglobulin; LRR: Leucine Rich Repeat; TLR: Toll/Interleukin-like Receptor; IL-1R: Interleukin-1 Receptor.

In keeping with past work^26,42^, we find that both the copy number of TLR, IL-1R and TIR-only genes, and the total abundance of their component TIR, LRR and Ig domains varies greatly across coral genomes (**Table 1**).

**Table 1.**
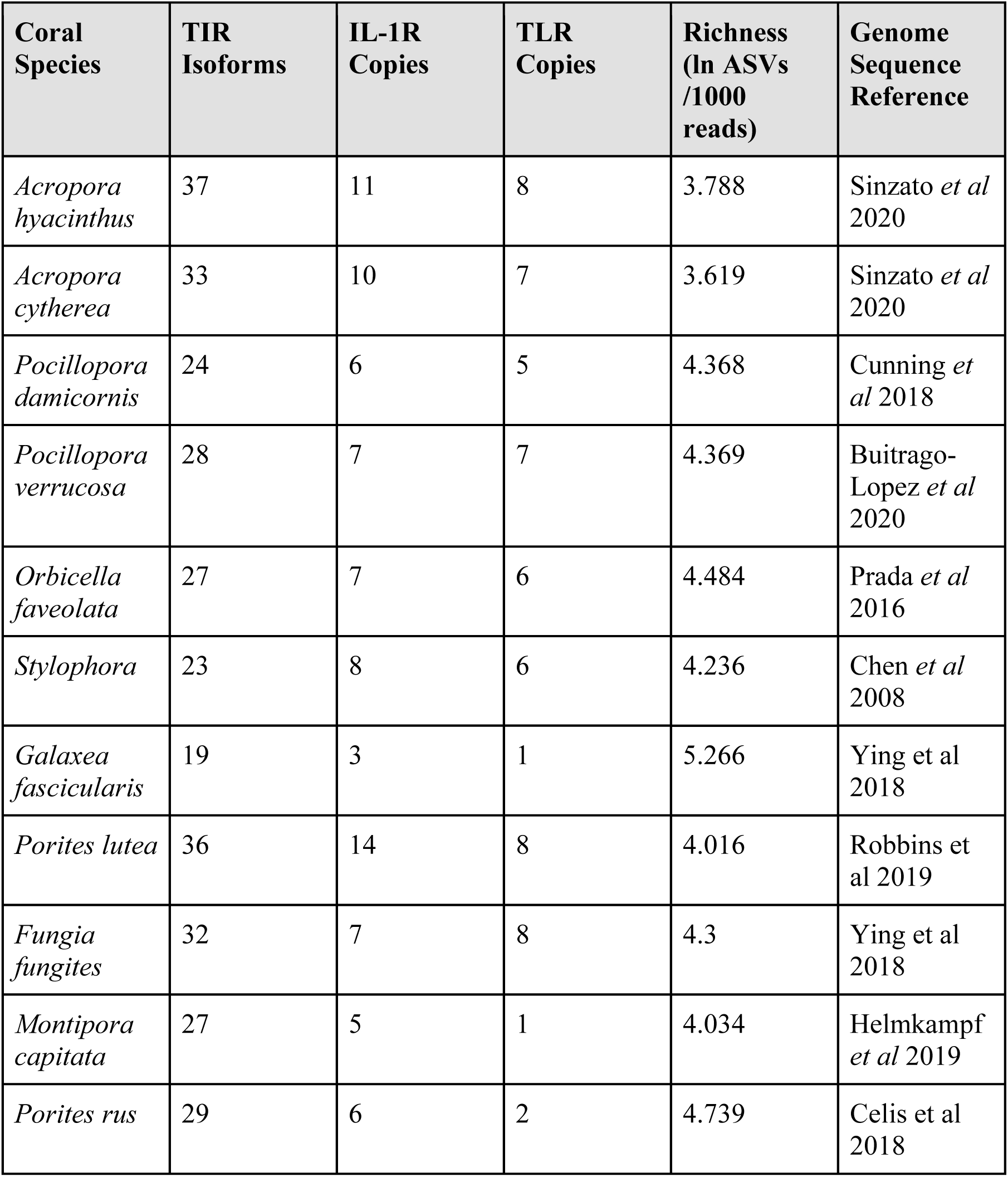
TIR domain containing elements and their associated numbers per species

### Coral IL-1R and TLR gene family copy number correlate with overall microbiome richness and evenness

In order to determine whether TLR or IL-1R might regulate microbiome biodiversity, we used phylogenetic independent contrasts (PIC) analysis to correlate changes in the genomic copy number of IL-1R and TLR against changes in microbiome richness or evenness during coral evolution.

We measured microbiome richness as the natural log of the observed number of amplicon sequence variants (ASVs) per 1000 sequences (**Table S5A**) (see Methods). Microbiome richness was significantly reduced by increases in the copy number of IL-1R (PIC R^2^ = 0.835, qFDR = 0.00055; **Fig. 2B, D**; **Fig. S1C, D**; **Table S6A**), or TLR (PIC R^2^ = 0.707, qFDR = 0.0076; **Fig. 2 C, D**; **Fig. S1E, F**; **Table S6A**). Thus, corals that harbor more TLR or IL-1R gene copies tend to have less diverse microbiomes, and vice versa. In manual inspection of the data, there were several striking examples of this statistical trend (**Table S7**). In the genus *Porites*, *P. lutea* had more than twice as many IL-1R and TLR gene copies as the closely related *P. rus* (14 IL-1R copies and eight TLR gene copies in *P. lutea* vs. six IL-1R copies and two TLR copies in *P. rus*; **Table 1**). Consistent with the idea that *P. lutea’*s expanded gene repertoire may play a role in microbiome filtering, *P. lutea’s* microbiome was roughly half as diverse as that of *P. rus* (56 ASVs/1000 reads vs. 114 ASVs/1000 reads, across all sample types).

**Fig. 2.**
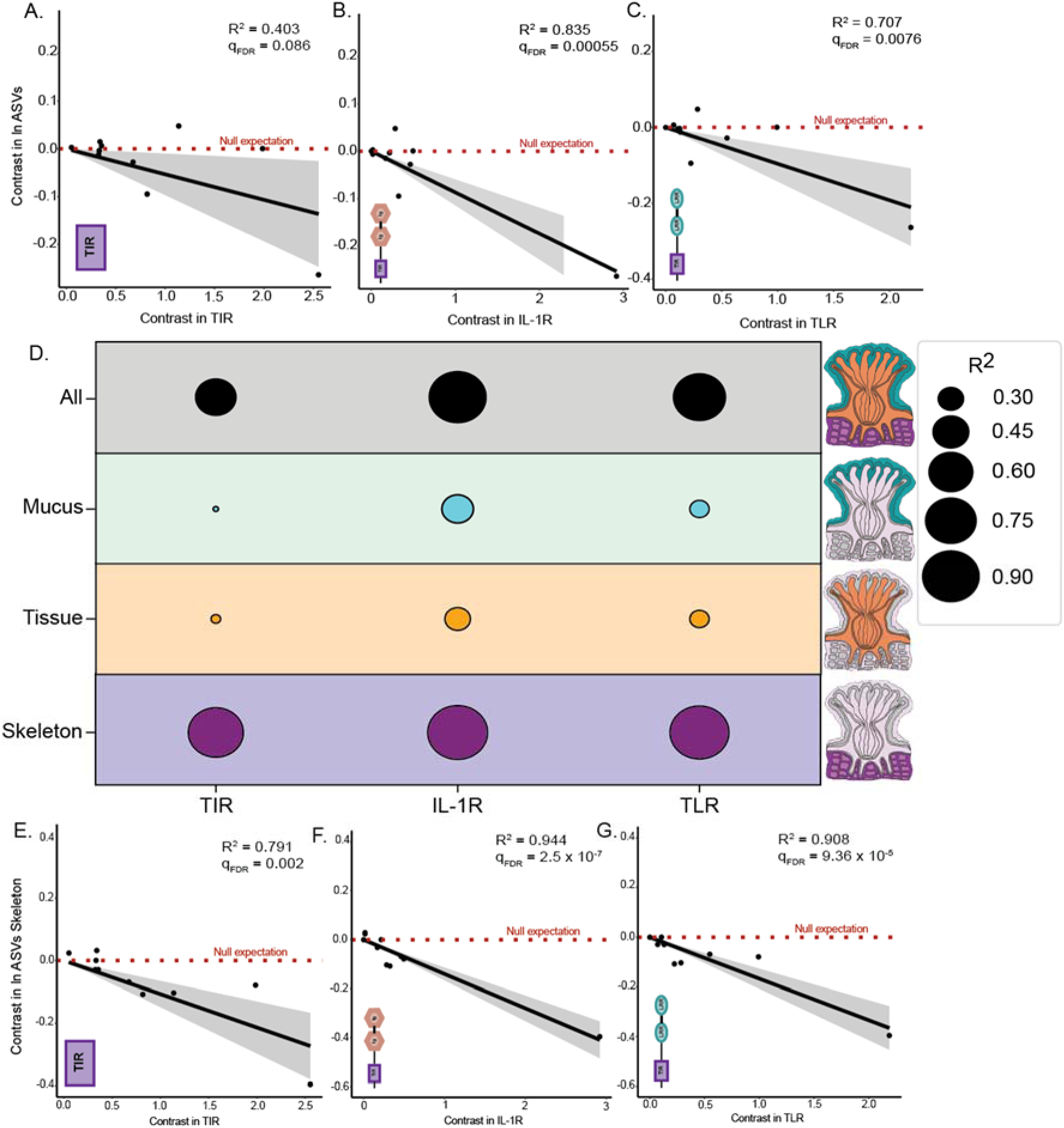
Rows show phylogenetic independent contrast analysis of genomic copy number of innate immune components against log microbiome richness (ln ASVs), for all **A.** TIR domains (R^2^ =0.403, qFDR = 0.086, p = 0.021); **B.** IL-1R genes (R^2^ = 0.835, qFDR = 0.00055, p = 5.13 x 10^-15^); **C.**TLR genes (R^2^ = 0.707, qFDR = 0.0076, p = 0.0012). Shading in phylogenetic independent contrasts analysis indicates the 95% confidence interval of the mean. **D.** R^2^ values from correlations of the genomic copy number of innate immune components against microbiome richness, organized by compartment. The bottom row uses microbiome data from coral skeleton only to show phylogenetic independent contrasts of innate immune components vs. log microbiome richness for: **E.** TIR-only genes (R^2^ =0.791, qFDR=0.002, p = 0.000252); **F.** IL-1R genes (R^2^ = 0.944, qFDR=2.15 x 10^-5^, p = 6.39 x 10^-7^); **G.** TLR genes (R^2^ = 0.90798, qFDR= 9.36 x 10^-5^, p = 5.85 x 10^-6^). No correlations had outliers, as defined by studentized residuals of >3 in absolute value.

We also quantified the microbiome evenness of each coral species in the analysis using the Gini Index (**Table S5B**), which takes on its highest value when ecological communities are least even. Gini index scores were strongly and significantly negatively correlated with the genomic copy number of IL-1R genes (PIC R^2^ = 0.876, qFDR = 0.000357; **Fig. S5C, D**; **Table S6A**) and TLR genes (PIC R^2^ = 0.805, qFDR = 0.0014; **Fig. S5E, F**; **Table S6A**), indicating that corals with more IL-1R or TLR gene copies have higher evenness.

These results support the prior hypothesis that IL-1R and TLR gene family expansions may influence coral microbiome structure. The magnitude of the effect was, however, quite surprising, with IL-1R gene copy number explaining ∼83% of the variation in microbiome richness, and ∼88% of the variation in microbiome evenness, among coral species in the analysis. Although necessarily limited to coral species for which genomes are available, these correlations are much stronger than the effects of several biotic and abiotic factors in the same GCMP dataset, including depth, temperature, and turf-algal contact^3^.

### Endolithic skeleton microbiomes drive correlations between TLR and IL-1R gene family expansion and microbiome structure

To this point, all our results were conducted by correlating genomic features of corals against the overall microbiome diversity of all available microbiome samples from each species in the GCMP dataset. However, microbiome richness has been shown to vary between coral compartments^3^. We expected that host tissues — where the immune system could most obviously act — would have microbiomes that most closely correlate with the innate immune gene repertoire of the host.

To test this idea, we separated coral microbiome samples into those that derive from mucus, tissue or endolithic skeleton, and repeated tests for correlations between coral innate immune repertoires and microbiome richness and evenness within each of those regions of anatomy.

Contrary to our hypothesis, coral endolithic skeleton — not tissue or mucus — was the sole driver of correlations between coral innate immune repertoire and microbiome richness (**Fig. 2E-G**; **Fig. S1-4**; **Table S6, 8**) and evenness (**Fig. S5-8**; **Table S6, 8**). Microbiome richness in coral endolithic skeleton was significantly correlated with IL-1R (R^2^ = 0.944, qFDR = 2.05 x 10^-5^; **Fig. 2F**, **Fig. S4C, D**; **Table S6D**), and TLR (R^2^ = 0.90798, qFDR = 9.36 x 10^-5^; **Fig. 2G**; **Fig. S4E, F**; **Table S6D**) gene copies, whereas tissue (**Fig. S3**; **Table S5C**) and mucus (**Fig. S2**; **Table S6B**) microbiome richness was not. Microbiome evenness showed similar patterns (**Fig. S6-8**; **Table S6, 8**).

Previous studies have reported that coral endolithic skeleton microbiomes are more species-rich than mucus or tissue, and show both species-specificity and signals of phylosymbiosis with their coral hosts^3^. Our results further suggest that endolithic skeleton microbiome diversity has tracked gene family expansions or contractions of coral TLR and IL-1R genes over evolution.

### Microbiome composition varies with IL-R and TLR gene copy numbers

In addition to regulating microbiome richness and evenness, coral innate immune systems may also influence coral microbiome composition. If so, we might expect microbiome composition to correlate with the repertoire of innate immune proteins encoded in coral genomes. To test this, we compared differences in overall microbiome composition for each pair of coral species using two phylogenetic beta diversity metrics: Weighted UniFrac and Unweighted UniFrac. Using these beta-diversity distance metrics, we conducted principal coordinates analysis (PCoA), using the first three PC axes (PC1,PC2, and PC3) of the PCoA ordination (Methods). We correlated the microbiome PC coordinates against the number of predicted TLR or IL-1R gene copies (**Fig. 3**; **Table S9**).

**Fig. 3.**
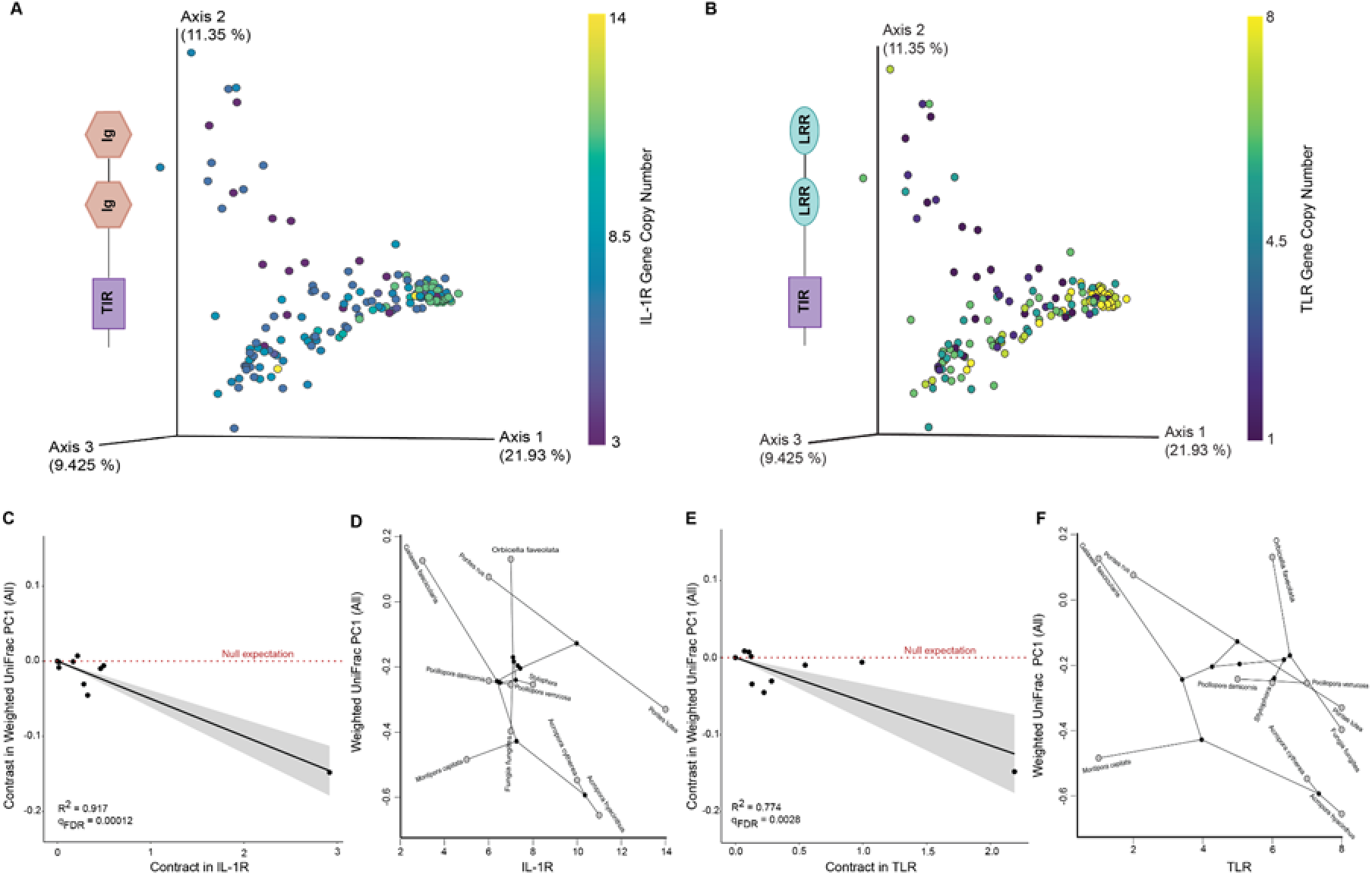
Comparison of innate immune gene repertoire and microbiome composition. **A.** Principle Coordinates Analysis (PCoA) of Weighted UniFrac Beta diversity distances for samples from all coral compartments, colored by genomic IL-1R copy number in the coral host. **B.** PCoA ordination of Weighted UniFrac distances for all compartments colored by genomic copy number for TLR genes. **C.** Phylogenetic independent contrasts regression of IL-1R copy number vs. Weighted UniFrac PC1 (R^2^ = 0.917, qFDR = 0.00012). **D.** Phylomorphospace of IL-1R copy number vs. Weighted UniFrac PC1. **E**. Phylogenetic independent contrasts regression of TLR copy number vs. Weighted UniFrac PC1 (R^2^ = 0.774, qFDR = 0.0028). **F.** Phylomorphospace of IL-1R copy number vs. Weighted UniFrac PC1. No correlations had outliers, as defined by studentized residuals of >3 in absolute value.

Innate immune gene copy numbers strongly and significantly correlated with microbiome composition. When all samples were considered together (irrespective of anatomical compartment) Weighted UniFrac PC1 correlated strongly with the genomic copy number of IL-1R genes (PIC Weighted UniFrac PC1 R^2^ = 0.917, qFDR = 0.00012; **Fig. 3A, C, D**; **Table S9A**) and TLR genes (PIC Weighted UniFrac PC1 R^2^ = 0.77, qFDR = 0.0028; **Fig. 3B, E, F**; **Table S9A**). Unweighted UniFrac PC1 showed similar trends (**Table S9A**). These results indicate that gene family expansions or contractions of the TLR and IL-1R gene families over evolution corresponded to dramatic changes in overall microbiome composition.

We repeated the above protocol separately on the mucus, tissue, and skeleton compartments separately (**Table S9B-D**). In coral mucus, tissue and skeleton, IL-1R gene copy number significantly correlated with Weighted UniFrac PC1 of the microbiome (PIC R^2^ = 0.72-0.81, FDR q < 0.01; **Table S9A-C**). TLR gene copy number significantly correlated with Weighted UniFrac PC1 in coral mucus and tissue (PIC R^2^ = 0.73, qFDR = 0.004; **Table S9B, C**) but not endolithic skeleton (**Table S9D**). In qualitative (presence/absence) analysis of microbiome β-diversity, IL-1R gene copy number correlated with Unweighted UniFrac PC1 in coral tissue and skeleton compartments, but not mucus (**Table S9B-D**), while TLR gene copy number correlated with Unweighted UniFrac PC1 in coral tissue but not mucus or skeleton (**Table S9B-D**).

These β-diversity results were more complex than the straightforward pattern of associations between innate immune gene copy number and microbiome richness. In coral tissue both IL-1R gene copy number and TLR gene copy number correlated with key aspects of quantitative (Weighted UniFrac PC1) and qualitative (Unweighted UniFrac PC1) microbiome β-diversity, in keeping with our original expectation that immunity should act strongly on the tissue-associated microbiome. In mucus, both TLR and IL-1R gene copy numbers appeared correlated with microbiome β-diversity when using quantitative but not qualitative metrics, suggesting immunity might influence microbial relative abundance more strongly than microbial presence/absence. In skeleton, IL-1R appeared correlated with IL-1R gene copy number, but not TLR gene copy number regardless of β-diversity metric. Together, these results suggest anatomically specific correlations between gene family expansion of some key innate immune genes and microbiome β-diversity.

### IL-1R and TLR copy number is associated with differential abundance of key microbes

We sought to identify microbial taxa that may be influenced by IL-1R or TLR gene copy number. This analysis could be confounded by the compositional nature of coral microbiome data. Therefore ANCOM-BC^41^, which accounts for compositionality, was used for the analysis. We first tested whether IL-1R copy number correlated with microbial differential abundance in all samples regardless of tissue compartment (i.e. ‘all’). In this overall analysis, IL-1R copy number correlated with the differential abundance of 102 families of bacteria and archaea, while TLR copy number correlated with 88 families (**Table S10A**).

Correlations between microbiome composition and IL-1R or TLR copy number varied with anatomy. The abundance of 38 bacterial families were significantly correlated with TLR copy number (ANCOM-BC qFDR < 0.05 in mucus, tissue, and skeleton; **Tables S10B-D**; **Fig. S11-S13**) and 38 bacterial families were also significantly and consistently correlated with IL-1R copy number (ANCOM-BC qFDR < 0.05 in mucus, tissue, and skeleton; **Table S10B-D**; raw heatmaps in **Fig. S14-S16**), both regardless of compartment. In contrast to the bacterial families with consistent interrelationships, some were specific to one or more compartments. One example are Rickettsiaceae, which live intracellularly in host cells. Rickettsiaceae relative abundance in tissue decreased with IL-1R (ANCOM-BC coef = −0.011, W = 0.33, qFDR = 0; **Fig. 4**, **Fig. S15**; **Table S10C**) and TLR (ANCOM-BC coef = −0.028, W =1.36, qFDR = 0; **Fig. 4**, **Fig S12**, **Table S10C**), but this taxon was not present in mucus (**Fig. S11**, **Fig S14**) or skeleton (**Fig. S13**, **Fig. S16**). *Nitrosopumiliaceae* archaea were notable for correlating with IL-1R and TLR gene copy number across all compartments (all ANCOM-BC qFDR < 0.05; **Fig S11-S16**).

**Fig. 4.**
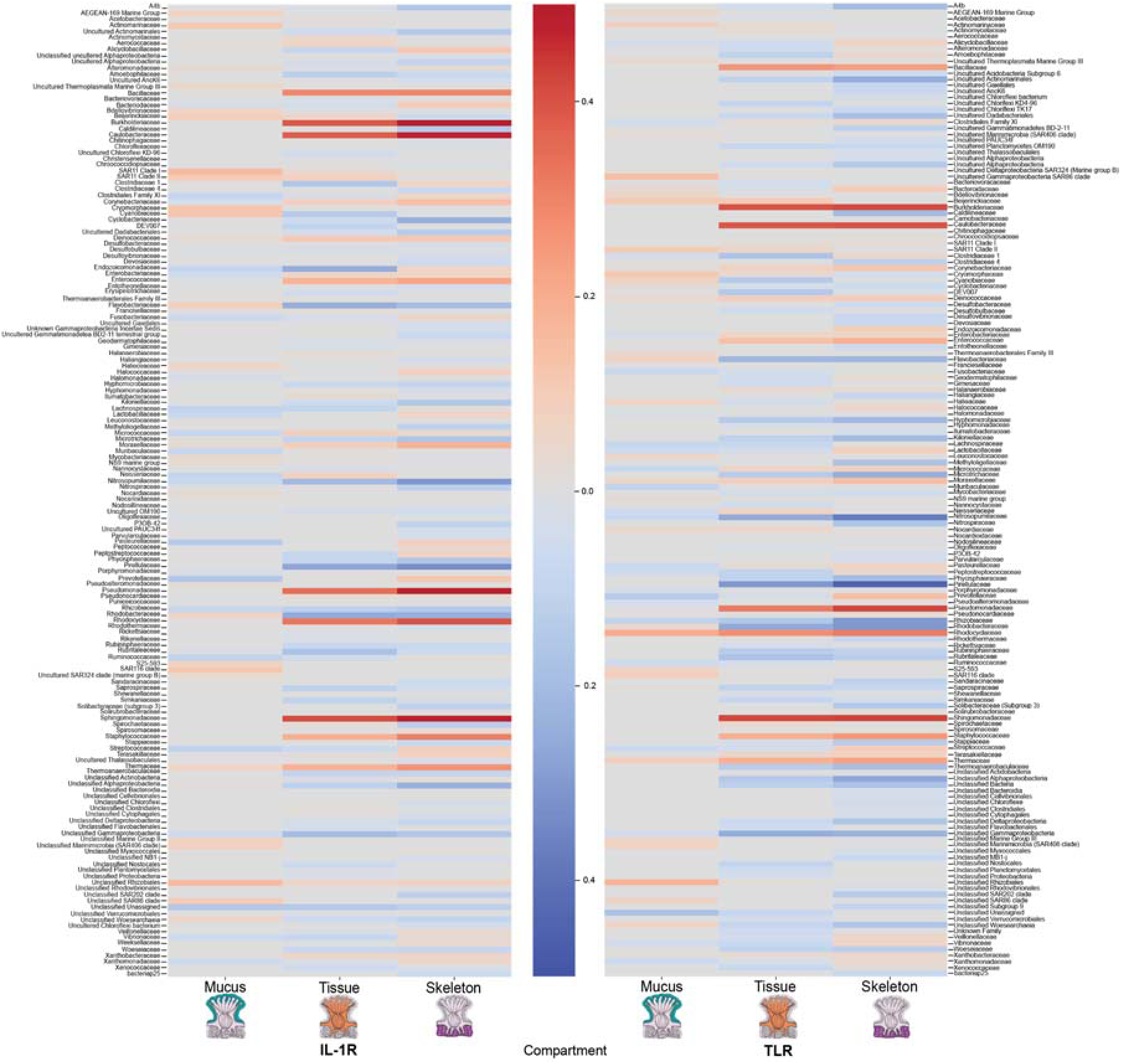
Heatmap showing microbial families significantly correlated with IL-1R and TLR copy numbers. Columns show correlations between the relative abundance of microbial families and IL-1R and TLR copy numbers in the mucus, tissue, and skeleton, respectively.

### The domain architecture of TLR but not IL-1R genes is associated with microbiome richness

IL-1R and TLR genes are known to vary in the domain architecture of their extracellular regions, with variable numbers of Ig or LRR sensing domains, respectively. We reasoned that if higher copy numbers of TLR or IL-1R genes are associated with microbiome richness because they act as filters on the microbiome, the same selective pressures (for greater specificity in microbial associates) might also influence the domain architecture of extracellular sensing components within TLR or IL-1R genes. To test this, we analyzed the average number of LRR and Ig domain copy numbers associated with each TLR or IL-1R gene in each coral species.

Interestingly, we find that there is a significant positive correlation between LRR domain copy number per TLR genes and TLR copies (R^2^ = 0.437, p = 0.0159; **Fig. 5A, B**; **Fig. S9A**) but not Ig copy number with IL-1R (R^2^ = 0.00067, p = 0.345; **Fig. 5C, D**; **Fig. S9B**). In other words, corals with more TLR genes also have TLR genes with more LRR domains each.

**Fig. 5.**
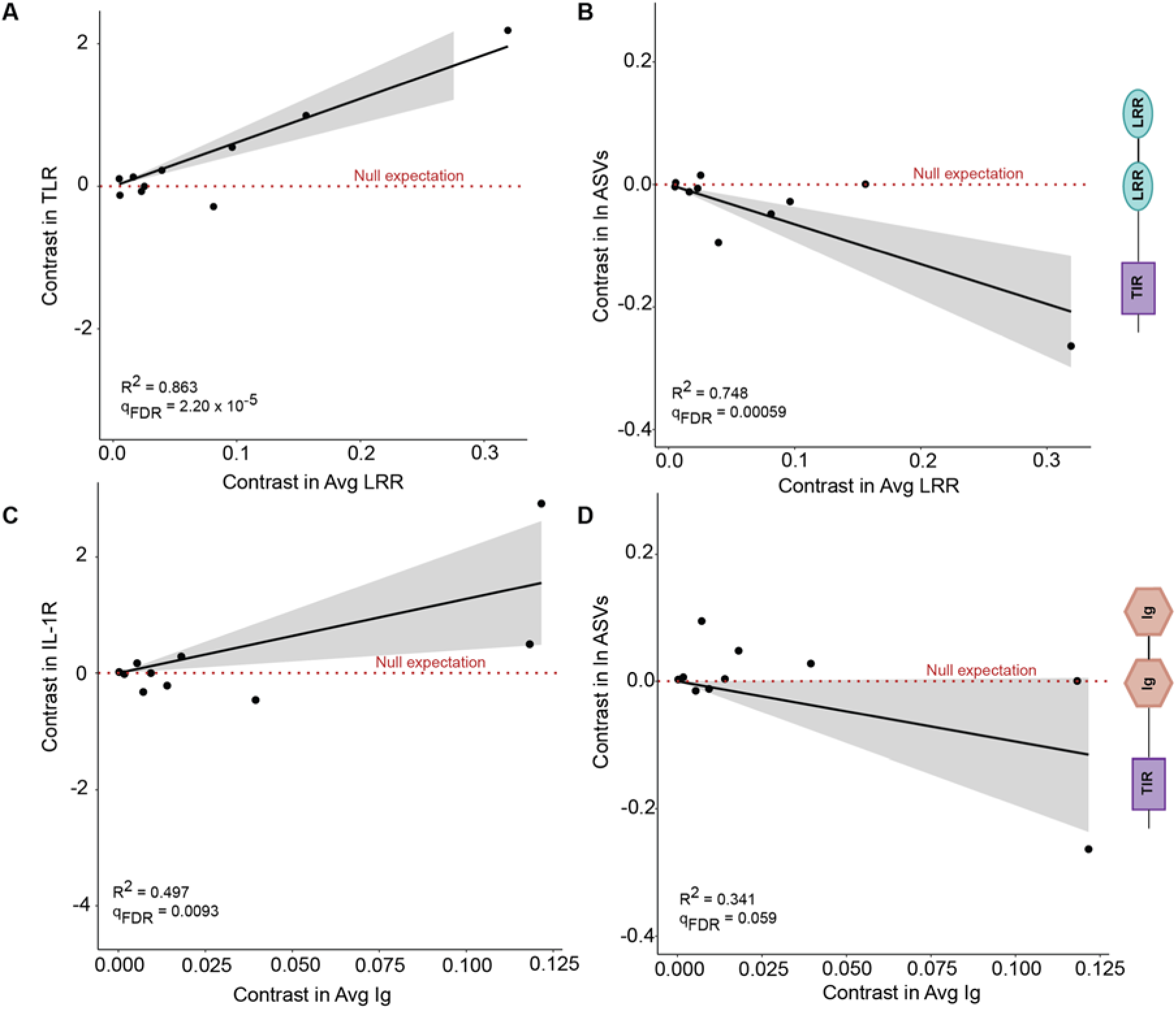
Domain copy numbers influence microbiome richness. Phylogenetic contrasts comparing **A.** average LRR domain copies per TLR vs TLR domain copy number (R^2^ = 0.863, qFDR = 2.20 x 10^-5^, **B.** average LRR domain copies per TLR vs ln ASVs (R^2^ = 0.748, qFDR = 0.00059), **C.** average Ig domain copies per IL-1R vs IL-1R copy number (R^2^ = 0.497, qFDR = 0.0093), and **D.** average Ig domain copies per IL-1R vs ln ASVs per genome (R^2^ = 0.341, qFDR = 0.059).

Furthermore, we find that microbiome richness is correlated negatively with the average number of LRR domain copies (PIC R^2^ = 0.748, p = 0.00059; **Fig 5B**) but not Ig domain copies (PIC R^2^ = 0.341, p = 0.059; Fig 5D). That is, corals whose TLR genes typically have more elaborate domain architectures also tend to have simpler (or more specific) microbial associations. Importantly, the genomic copy number of LRR or Ig domains in total (rather than per gene) did not correlate with microbiome richness and beta diversity (**Table S8, S11**), so this finding does not simply recapitulate our previous findings regarding gene copy number.

### Responses to IL-1R and TLR are correlated between bacterial families

The microbe associated molecular patterns (MAMPs) detected by species-specific members of the TLR and IL-1R gene families could in principle be the same or independent. If independent, the correlation between any given microorganism and TLR gene family expansions would be independent of IL-1R gene family expansions. To test this, we analyze whether the correlation coefficients between each microbial family’s relative abundance with TLR or IL-1R copy number were themselves correlated with one another. Microbial families’ responses to IL-1R and TLR gene family copy numbers were themselves very strongly correlated (R^2^ = 0.9062; **Fig S11**). Thus, microbes that are in high abundance in corals with many IL-1R gene copies also tend to be in high abundance in corals with many TLR gene copies. This suggests that TLR and IL-1R gene family expansions sculpt coral microbiomes in similar, rather than contrasting ways.

## Discussion

By combining coral microbiome and genomic data in a comparative framework, we demonstrate that TIR-domain containing innate immune gene repertoires strongly correlate with microbiome structure and diversity. These results suggest that gene family expansions of innate immune genes may have contributed to differences in the structure of coral microbiomes across millions of years of evolution. Those differences in immunity and microbiome structure, in turn, may influence the ability of modern scleractinian corals to survive escalating challenges from disease and climate change.

### Coral innate immune systems may be filtering microbiome membership

A key finding of the Earth Microbiome Project was that the diverse communities of microbes associated with animals and plants are nonetheless much less diverse than most environmental microbiomes^28^. Indeed, these animal-associated microbiomes were found to show high nestedness relative to the environment, indicating that what lives on animals is mostly — though certainly not entirely — a subset of environmental microbes^28^. This suggests that filtering of environmental microbes is one key process shaping animal and plant microbiomes. That filtering is likely to be genomically encoded, since more closely related animals tend to have more similar microbiomes (‘phylosymbiosis’^3, 43–45)^, excepting important deviations from this trend such as those driven by the evolution of specialized diets^46,47^ or flight^48^. Could evolutionary changes in host immunity drive this trend, by altering which environmental microbes are excluded?

Our results in corals suggest that, for this group at least, immunity plays an important role in overall microbiome richness and evenness. Both coral microbiome composition^3^, and coral innate immune responses^49,50^ have long been known to vary between species. Within coral populations, there is evidence that variation in immune activity between coral fragments correlates with their microbiome structure. For example, in *Montipora capitata*, increases in phenoloxidase activity correlated with decreased microbiome richness^51^. If such correlations extended across species, it could begin to explain why coral microbiomes differ so greatly in bacterial and archaeal biodiversity. TIR-domain containing genes have undergone large gene-family expansions in some lineages of corals, which has been proposed as a driver of differences in coral microbiome richness^26^. Our findings support this hypothesis, and suggest that expansions of key innate immune genes reduce microbiome richness by allowing for sensing and exclusion of more diverse groups of microbes.

### Microbiome composition changes with TLR and IL-1R copy number

We find a relationship between microbiome β-diversity and the copy number of TIR-containing innate immune genes. Immune genes of animals and plants have in the past been reported to alter the β-diversity of symbiotic microbes, including commensals. For example, using Toll-like receptor (TLR) 5 deficient compared to wild type mice, TLR genes have been correlated with membership and host-microbial interactions with specific members of the mouse gut microbiome, although the overall effect of TLR on gut microbiome β-diversity remains controversial^52^. Similarly, *Arabidopsis* with loss of function mutations in the pattern recognition receptor (PRR) gene FLS2, have altered rhizosphere microbiome β-diversity relative to wild type controls^53^. Our findings extend these observations to coral microbiomes, and suggest that differences in innate immunity may explain substantial portions of the known differences in microbiome structure between coral species.

In our results, the relative abundances of diverse microbial taxa correlated with TLR or IL-1R gene copy number. Interestingly, these effects were not independent: microbes that correlated with TLR copy number also tended to be correlated with IL-1R copy number. This might reflect either that TLR and IL-1R gene family expansions are driven by similar selective pressures and therefore tend to co-occur, or that both types of gene family expansion influence the microbiome in similar ways.

### The relationship between innate immune gene repertoire and the microbiome is anatomically-specific

While there are clear reasons to expect coral innate immune gene repertoire to affect tissue-associated coral microbiomes, our results suggest that these gene family expansions have even more clear-cut effects on coral’s endolithic skeleton microbiomes. Corals show compartmentalized differences in molecular function^54^ and microbial richness and composition across anatomy^3^. Perhaps surprisingly, the microbiome of coral endolithic skeleton has been shown to be far more diverse than coral tissues or mucus^3^. In our results, endolithic skeleton microbiomes were more strongly correlated with host innate immunity than either mucus or tissue.

These results suggest that coral immunity strongly influences endolithic microbiomes. They also underline the importance of efforts to reevaluate how we conceptualize coral immunity and calcification. While coral calcification and immunity are typically thought of separately, recent work synthesizes these two fields^55^, noting that skeleton is a key barrier against pathogens, and in many species contains diverse defensive chemical compounds and enzymes. These notably include melanin, which is a key aspect of immune defense in many non-vertebrate lineages.

### Disease-susceptible corals have more, not fewer, TIR-domain containing genes

One counter-intuitive aspect of our results is that the coral taxa with the largest numbers of TIR-containing innate immune genes tend to be those, such as *Acropora,* regarded as more susceptible to disease^56^. This is surprising, because we might expect more disease-susceptible coral species to have less diverse innate immune gene repertoires. How gene-family expansion, the commensal microbiome, and coral disease interact is a rich topic for future investigation. Our results here suggest several hypotheses that could be explored. It could be that more diverse microbial ecosystems are less susceptible to invasion by pathogens, as per Elton’s 1958 biotic resistance hypothesis^57,58^. Alternatively, less rich microbiomes may be less functionally redundant, increasing the risk that loss of any particular beneficial microbe may degrade host health ^59,60^. Finally, the damage threshold hypothesis^61^ proposes that corals differ in immune strategy, with many slow-growing corals constitutively expressing a variety of innate immune genes, while other fast-growing corals adopt a reactive strategy, which avoids expending resources on baseline expression of many innate immune genes in order to maximize growth or fecundity. If so, then reactive, fast-growing corals may experience stronger selective pressures to diversify sensors that can detect pathogens or cellular damage, since their baseline level of protection is low. TIR-domain containing gene family expansions in these reactive corals may enable more sensitive and/or specific responses to pathogen exposure.

## Conclusions

We find that among sequenced coral genomes, gene family expansions of TLR and IL-1R genes are correlated with alterations in microbiome structure, and reductions in microbiome richness — with this apparent interplay between innate immunity and the microbiome most noticeable in coral endolithic skeleton. These findings are consistent with the idea that animal immunity sculpts microbiome structure and composition in part by sensing and filtering out many environmental microbes. This interpretation of the correlations we found between gene copy number and the microbiome is reinforced by the correlation between microbiome richness and the domain architecture within TLR genes, wherein there are more sensing (i.e. LRR) domains per TLR gene in corals that have lower microbiome richness. Our results further underscore the importance of distinguishing coral microbiomes across anatomy, and of exploring how coral innate immunity regulates corals’ diverse endolithic microbiomes. Finally, these results emphasize that integrating expanding coral genome and microbiome datasets in comparative frameworks is a promising approach that will help to uncover the interactions between immunity, the microbiome, and reef health.

## Data Availability

Biom and taxonomy tables are available on Qiita (Study ID 10895). Analysis code is available on GitHub: https://github.com/zaneveld/GCMP_Genomics/

## Acknowledgements

The authors would like to acknowledge many contributors for their field assistance during past collections of Global Coral Microbiome Project data reanalyzed in this manuscript, including: Rebecca Vega Thurber, a leader of the original GCMP project; Tasman Douglass, Margaux Hein, Frazer McGregor, Kathy Morrow, Katia Nicolet, Cathie Page, and Gergely Torda for their field assistance in Australia; Valeria Pizarro, Mateo López-Victoria, Styles Smith, Alaina Weinheimer, Claudia Tatiana Galindo for Assistance in Colombia; Chris Voolstra, Maren Ziegler, Anna Roik, and many others at KAUST for field assistance in Saudi Arabia; Mark Vermeij, Kristen Marhaver, Pedro Frade, Ben Mueller, and others at CARMABI for field assistance during sampling in Curaçao; Danwei Huang for field assistance in Singapore; Ruth Gates, Katie Barrott, Courtney Couch, Keoki Stender for field assistance in Hawai’i; Le Club de Plongee Suwan Macha and Jean-Pascal Quod for field assistance, and Amelia Foster and Jerome Payet for sampling in Isle de la Réunion; the Burkepile lab for field assistance in Mo’orea; Yossi Loya, Raz Tamir for sampling assistance in Israel; and Lyndsy Gazda, Jamie Lee Proffitt, Gabriele Swain, and Alaina Weinheimer for their assistance in the laboratory. The authors also acknowledge the staff of the Coral Bay Research Station, Lizard Island Research Station, Lord Howe Island Marine Park, Lord Howe Island Research Station, and RV Cape Ferguson for their logistical support. This work was supported by a The GCMP dataset was supported by a National Science Foundation Dimensions of Biodiversity grant (#1442306) to Rebecca Vega Thurber and MM, with sequencing provided by an in-kind UC San Diego Seed Grant in Microbiome Science grant to JZ. Analysis in this manuscript was supported by NSF IoS CAREER grant (# 1942647) and an in-kind UC San Diego Seed Grant in Microbiome Science grant to JZ. GCMP data collection was supported by a National Science Foundation Dimensions of Biodiversity grant (#1442306) to M.M.

## Author Contributions

TB, DS, JZ analyzed the data; TB, JZ, DS wrote the manuscript; RM, JFP, MM, and JZ collected GCMP data and metadata; all authors edited the manuscript.

## Supplementary Figures

Supplementary Figures 1-4: Richness vs. gene copy number for 1) All compartments 2) mucus, 3 tissue, 4 skeleton

Supplementary Figures 5-8: Evenness vs. gene copy number for 1) All compartments 2) mucus, 3 tissue, 4 skeleton

Supplementary Figure 9: Comparison of domain copy number within genes and gene copies within genomes.

Supplementary Figure 10: Correlated responses of microbial families to TLR and IL-1R gene copy number

Supplementary Figures 11-16. Relative abundance of microbial classes in mucus 11, tissue 12 and skeleton 13 samples sorted by TLR copy number or IL-1R copy number (14,15,16).

**Fig. S1.**
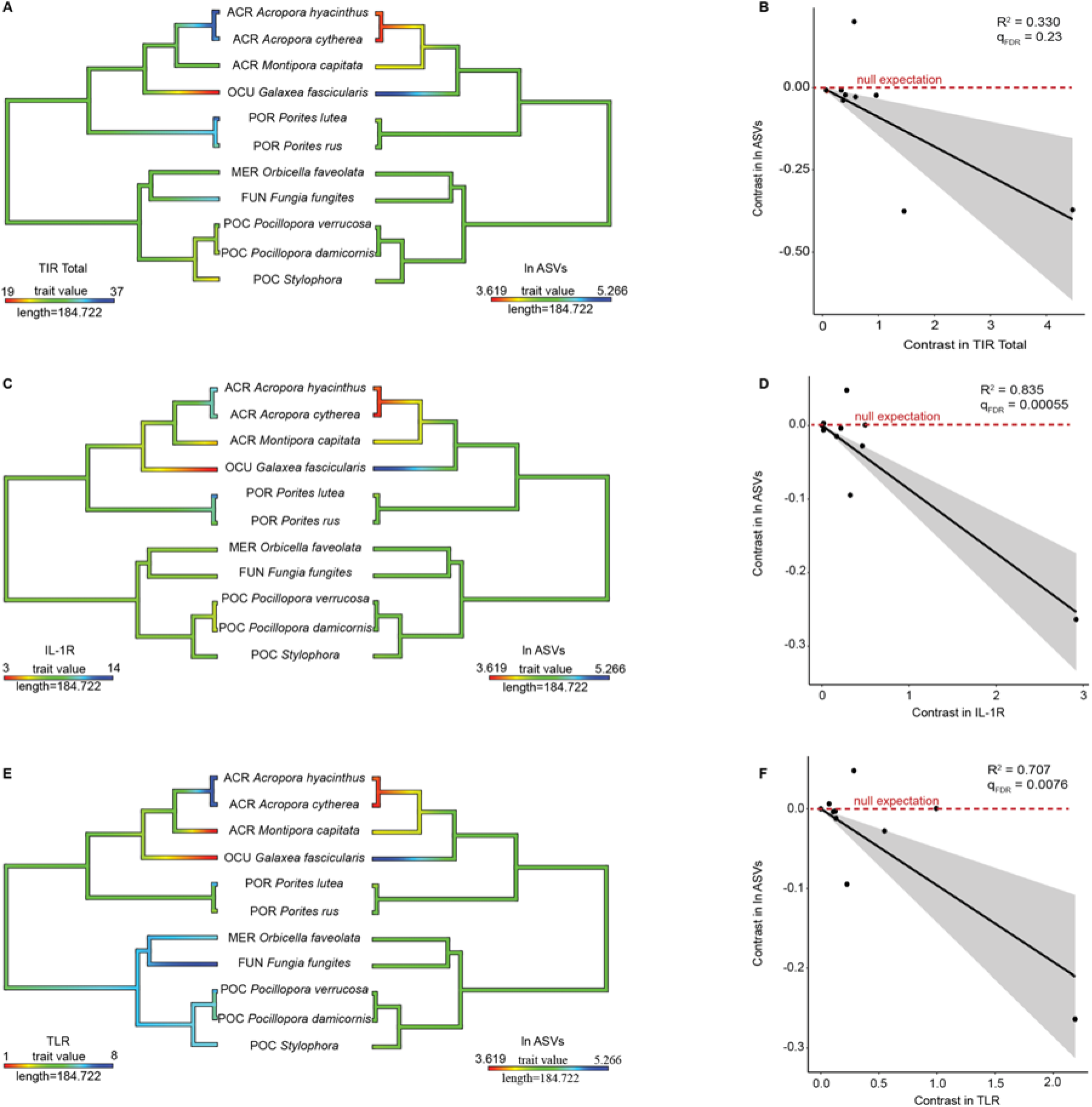
Phylogenetic comparison of coral microbiome richness and innate immune repertoire for all compartments combined. Rows show ancestral state reconstructions (left column) of innate immune gene copy number and microbiome richness, as well as phylogenetic independent contrast analysis for all predicted isoforms (right column) of **A,B** TIR-only genes (R^2^ =0.330, qFDR= 0.23,p = 0.065); **C,D** IL-1R genes (R^2^ = 0.834, qFDR= 0.00055, p = 5.14 x 10^-5^); **E,F** TLR genes (R^2^ = 0.707, qFDR=0.0076, p = 0.0012) compared against log microbiome richness (ln ASVs). Shading in phylogenetic independent contrasts analysis indicates the 95% confidence interval of the mean.

**Fig. S2.**
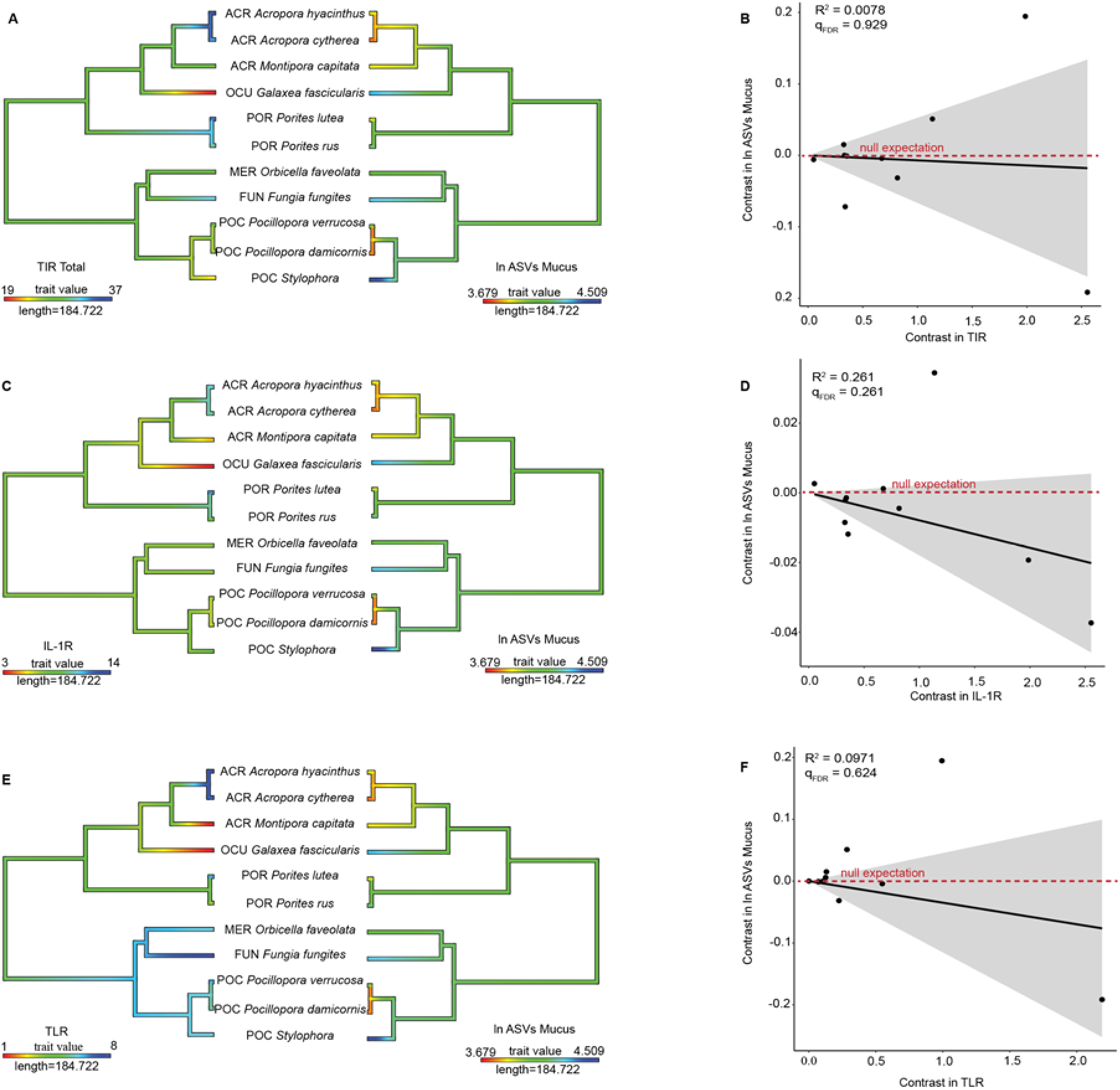
Phylogenetic comparison of coral microbiome richness and innate immune repertoire for the mucus compartment. Rows show ancestral state reconstructions (left column) of innate immune gene copy number and microbiome richness, as well as phylogenetic independent contrast analysis for all predicted isoforms (right column) of **A,B** TIR-only genes (R^2^ =0.0078, qFDR= 0.929,p = 0.796); **C,D** IL-1R genes (R^2^ = 0.261, qFDR= 0.261, p = 0.109); **E,F** TLR genes (R^2^ = 0.0971, qFDR=0.624, p = 0.351) compared against log microbiome richness (ln ASVs). Shading in phylogenetic independent contrasts analysis indicates the 95% confidence interval of the mean.

**Fig. S3.**
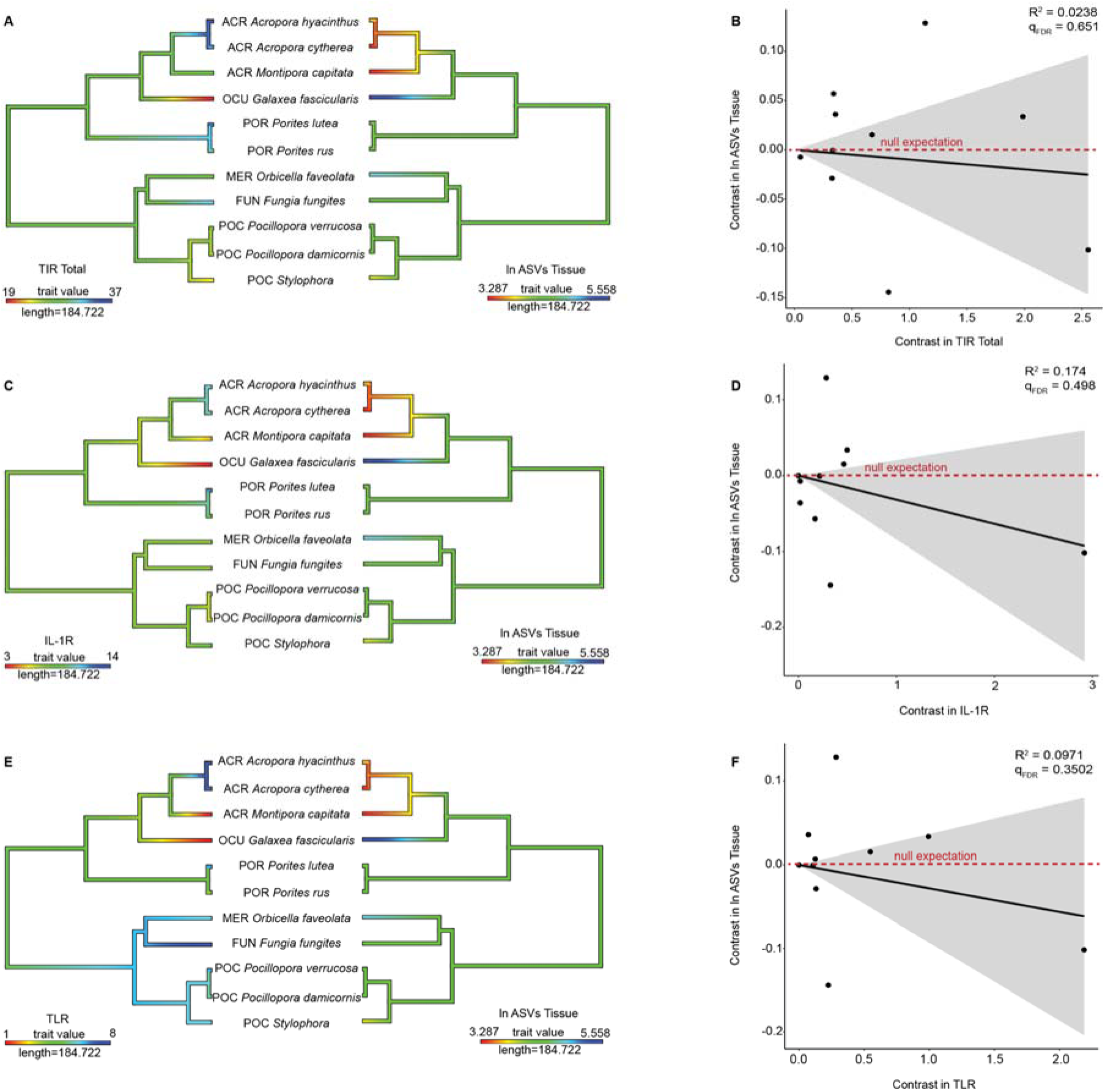
Phylogenetic comparison of coral microbiome richness and innate immune repertoire for the tissue compartment. Rows show ancestral state reconstructions (left column) of innate immune gene copy number and microbiome richness, as well as phylogenetic independent contrast analysis for all predicted isoforms (right column) of **A,B** TIR-only genes (R^2^ = 0.0238, qFDR=0.929, p = 0.651); **C,D** IL-1R genes (R^2^ = 0.174, qFDR=0.498, p = 0.202); **E,F** TLR genes (R^2^ = 0.0974, qFDR=0.624, p = 0.3502) compared against log microbiome richness (ln ASVs). Shading in phylogenetic independent contrasts analysis indicates the 95% confidence interval of the mean.

**Fig. S4.**
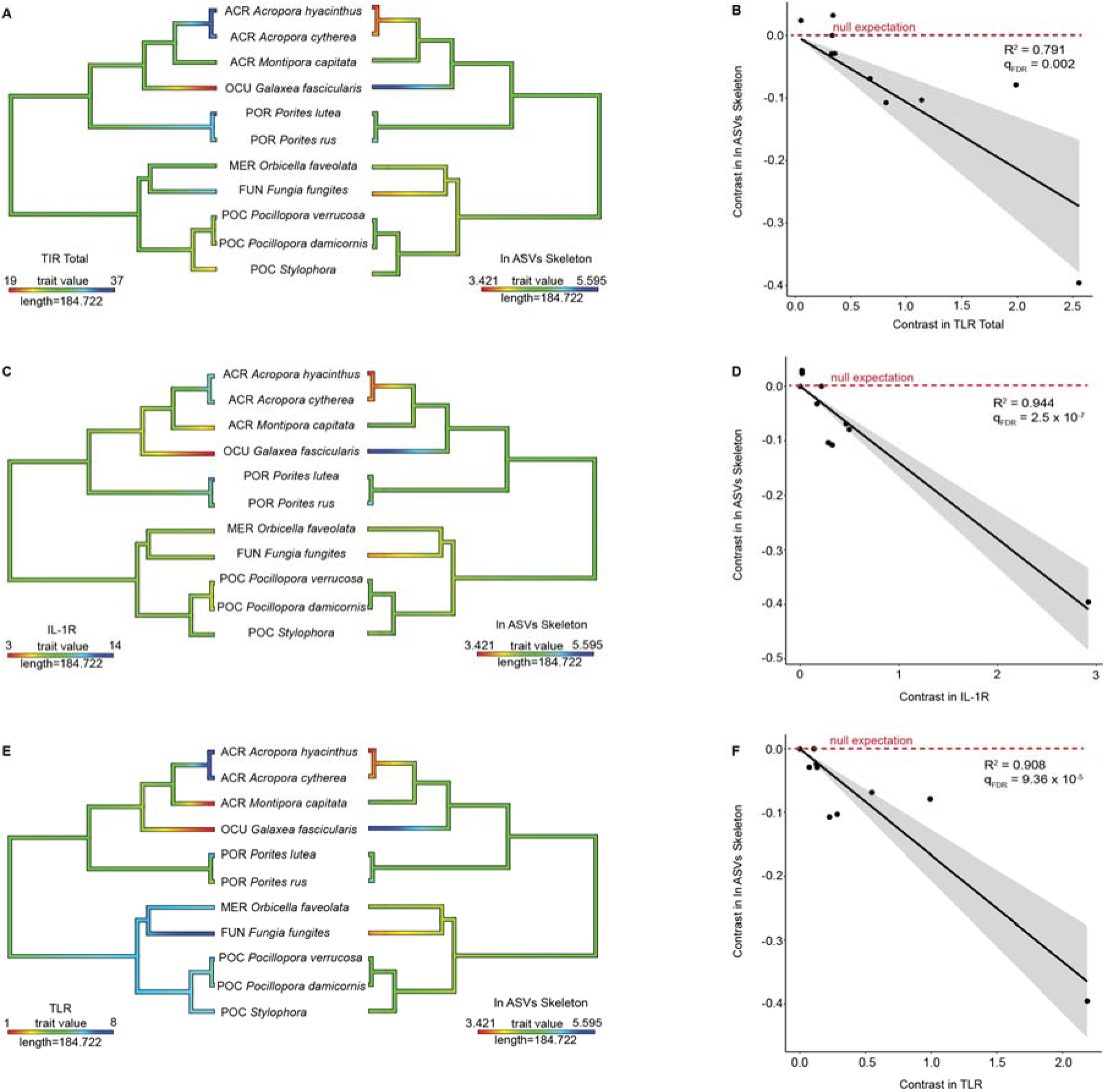
Phylogenetic comparison of coral microbiome richness and innate immune repertoire for the skeletal compartment. Rows show ancestral state reconstructions (left column) of innate immune gene copy number and microbiome richness, as well as phylogenetic independent contrast analysis for all predicted isoforms (right column) of **A,B** TIR-only genes (R^2^ =0.791, qFDR=0.002, p = 0.000252); **C,D** IL-1R genes (R^2^ = 0.944, qFDR=2.15 x 10^-5^, p = 6.39 x 10^-7^); **E,F** TLR genes (R^2^ = 0.90798, qFDR= 9.36 x 10^-5^, p = 5.85 x 10^-6^) compared against log microbiome richness (ln ASVs). Shading in phylogenetic independent contrasts analysis indicates the 95% confidence interval of the mean.

**Fig. S5.**
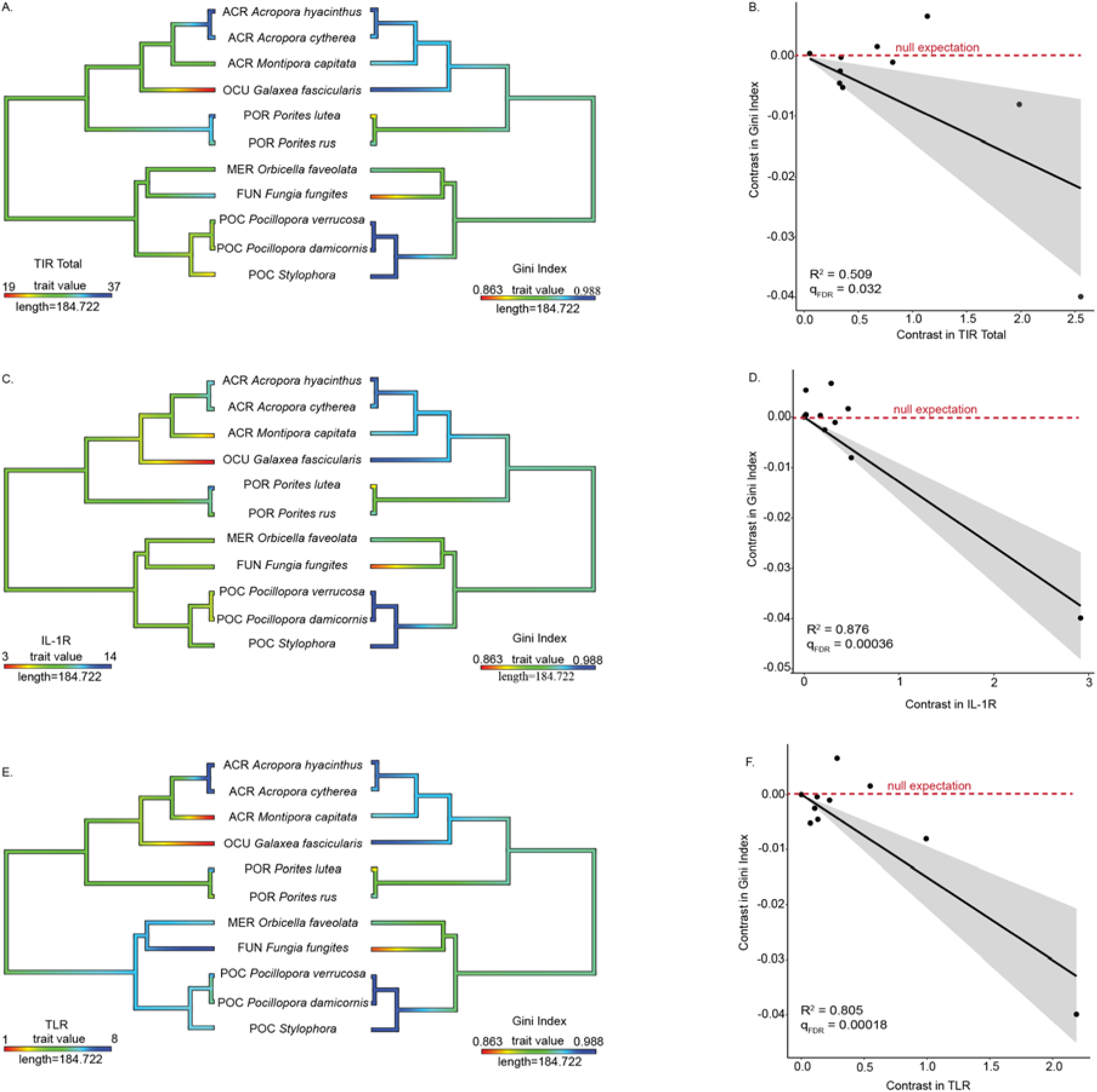
Phylogenetic comparison of coral microbiome evenness and innate immune repertoire in all compartments combined. Rows show ancestral state reconstructions (left column) of innate immune gene copy number and microbiome evenness (Gini index), as well as phylogenetic independent contrast analysis for all predicted isoforms (right column) of **A,B** TIR-only genes (R^2^ =0.509, qFDR=0.032, p = 0.0083); **C,D** IL-1R genes (R^2^ = 0.876, qFDR=0.00036, p = 2.30 x 10^-5^); **E,F** TLR genes (R^2^ = 0.805, qFDR= 0.0014, p =0.00018) compared against microbiome evenness (Gini index). Shading in phylogenetic independent contrasts analysis indicates the 95% confidence interval of the mean.

**Fig. S6.**
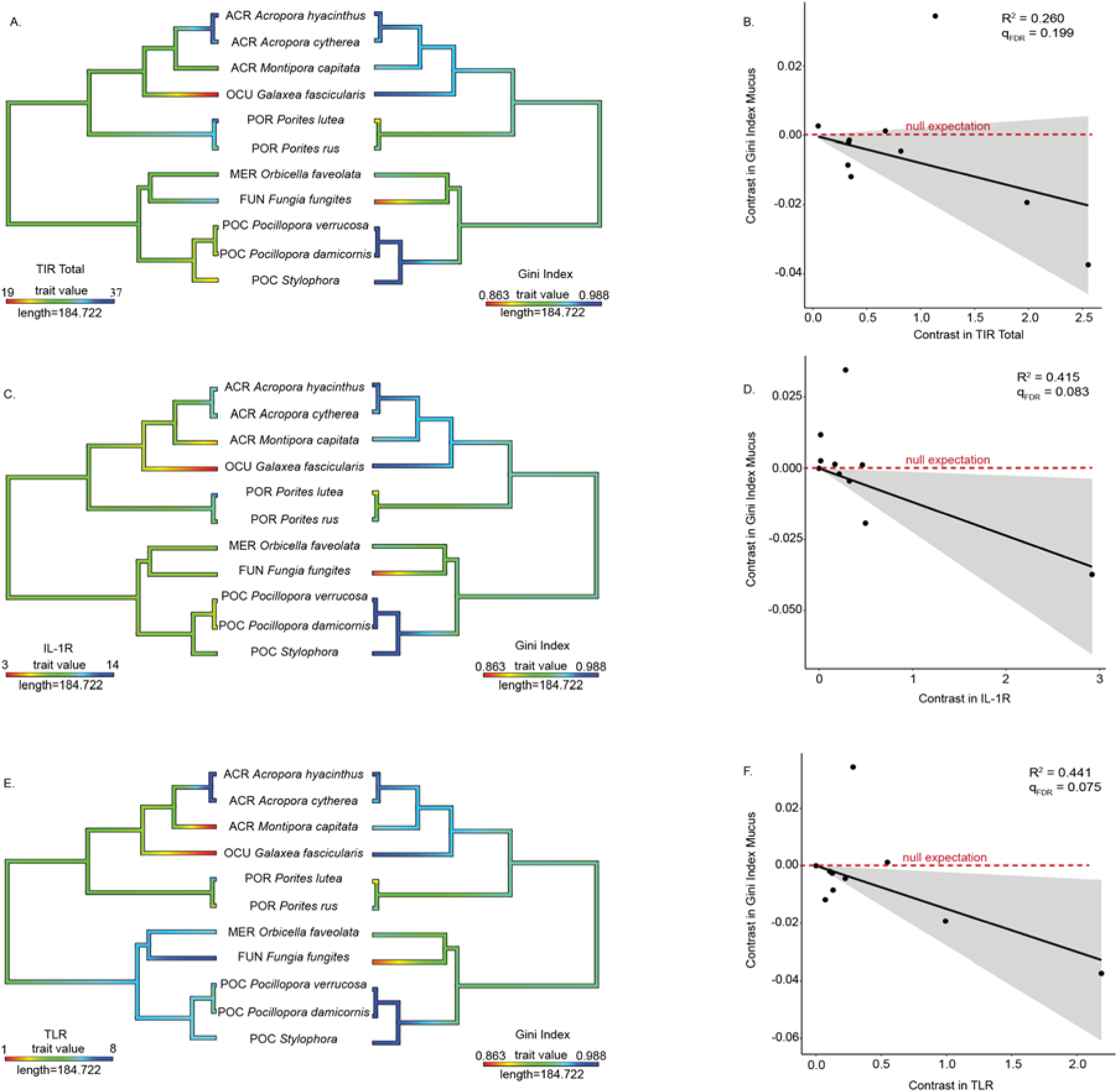
Phylogenetic comparison of coral microbiome evenness and innate immune repertoire in the mucus compartment. Rows show ancestral state reconstructions (left column) of innate immune gene copy number and microbiome evenness (Gini index), as well as phylogenetic independent contrast analysis for all predicted isoforms (right column) of **A,B** TIR-only genes (R^2^ =0.260, qFDR=0.199, p = 0.109); **C,D** IL-1R genes (R^2^ = 0.415, qFDR=0.083, p = 0.0323); **E,F** TLR genes (R^2^ = 0.441, qFDR= 0.075, p =0.0258) compared against microbiome evenness (Gini index). Shading in phylogenetic independent contrasts analysis indicates the 95% confidence interval of the mean.

**Fig. S7.**
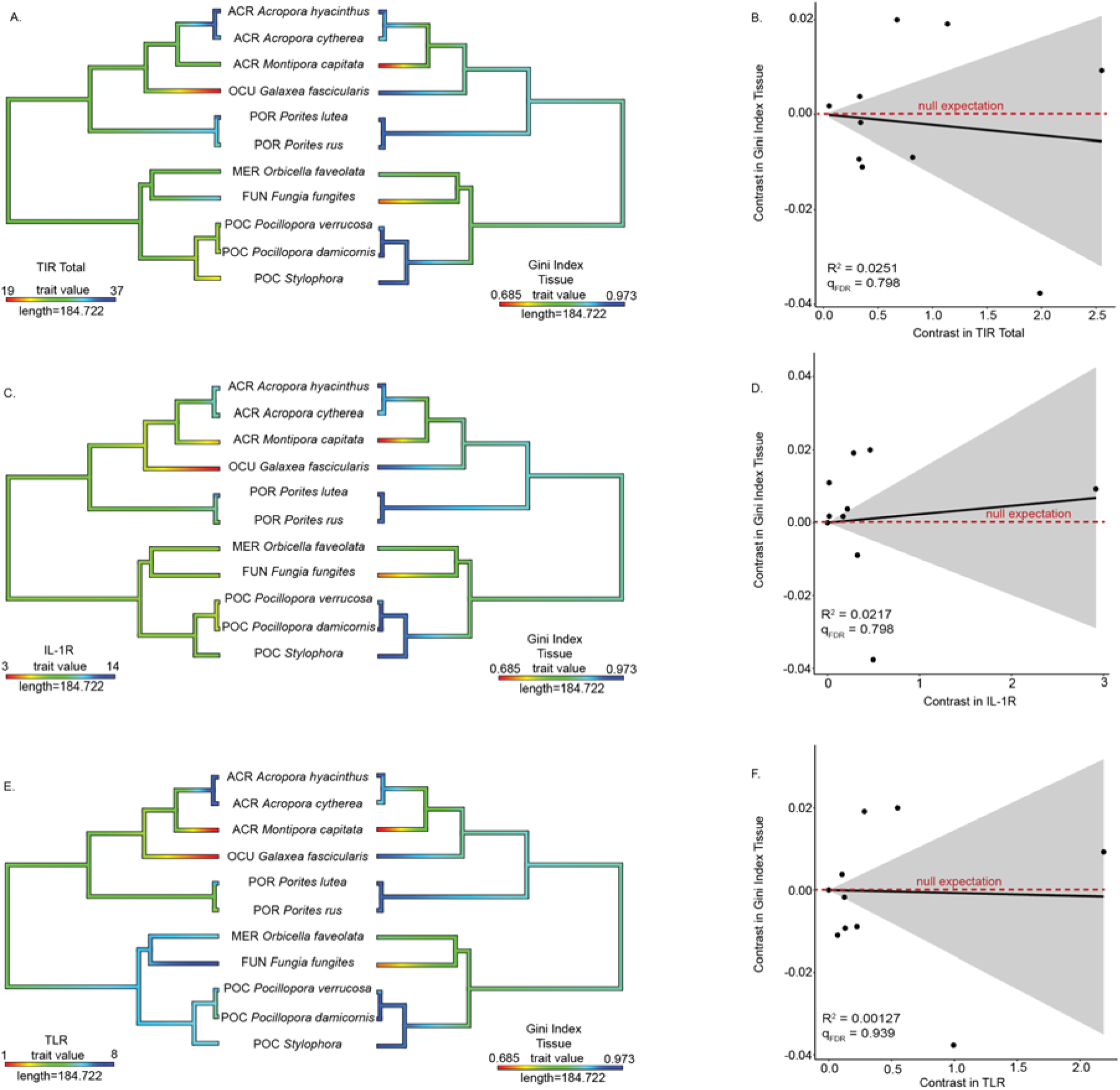
Phylogenetic comparison of coral microbiome evenness and innate immune repertoire in the tissue compartment. Rows show ancestral state reconstructions (left column) of innate immune gene copy number and microbiome evenness (Gini index), as well as phylogenetic independent contrast analysis for all predicted isoforms (right column) of **A,B** TIR-only genes (R^2^ =0.0251, qFDR=0.798, p = 0.6421); **C,D** IL-1R genes (R^2^ = 0.0201, qFDR=0.798, p = 0.678); **E,F** TLR genes (R^2^ = 0.00127, qFDR= 0.939, p =0.917) compared against microbiome evenness (Gini index). Shading in phylogenetic independent contrasts analysis indicates the 95% confidence interval of the mean.

**Fig. S8.**
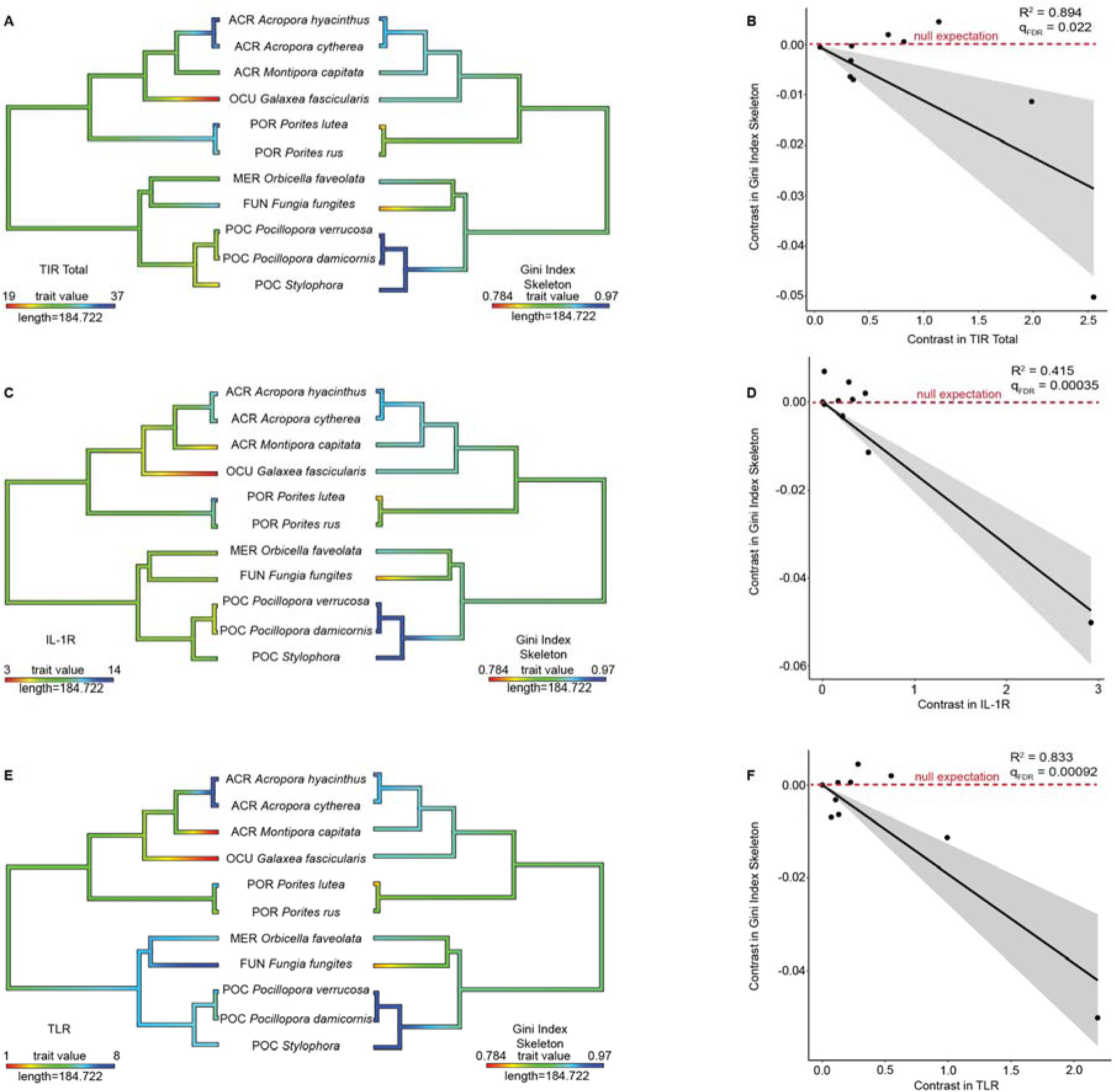
Phylogenetic comparison of coral microbiome evenness and innate immune repertoire in the skeleton compartment. Rows show ancestral state reconstructions (left column) of innate immune gene copy number and microbiome evenness (Gini index), as well as phylogenetic independent contrast analysis for all predicted isoforms (right column) of **A,B** TIR-only genes (R^2^ =0.894, qFDR=0.022, p = 1.12 x10^-5^); **C,D** IL-1R genes (R^2^ = 0.415, qFDR=0.00035, p = 0.0323); **E,F** TLR genes (R^2^ = 0.833, qFDR=0.00092, p =8.86 x 10^-5^) compared against microbiome evenness (Gini index). Shading in phylogenetic independent contrasts analysis indicates the 95% confidence interval of the mean.

**Fig. S9.**
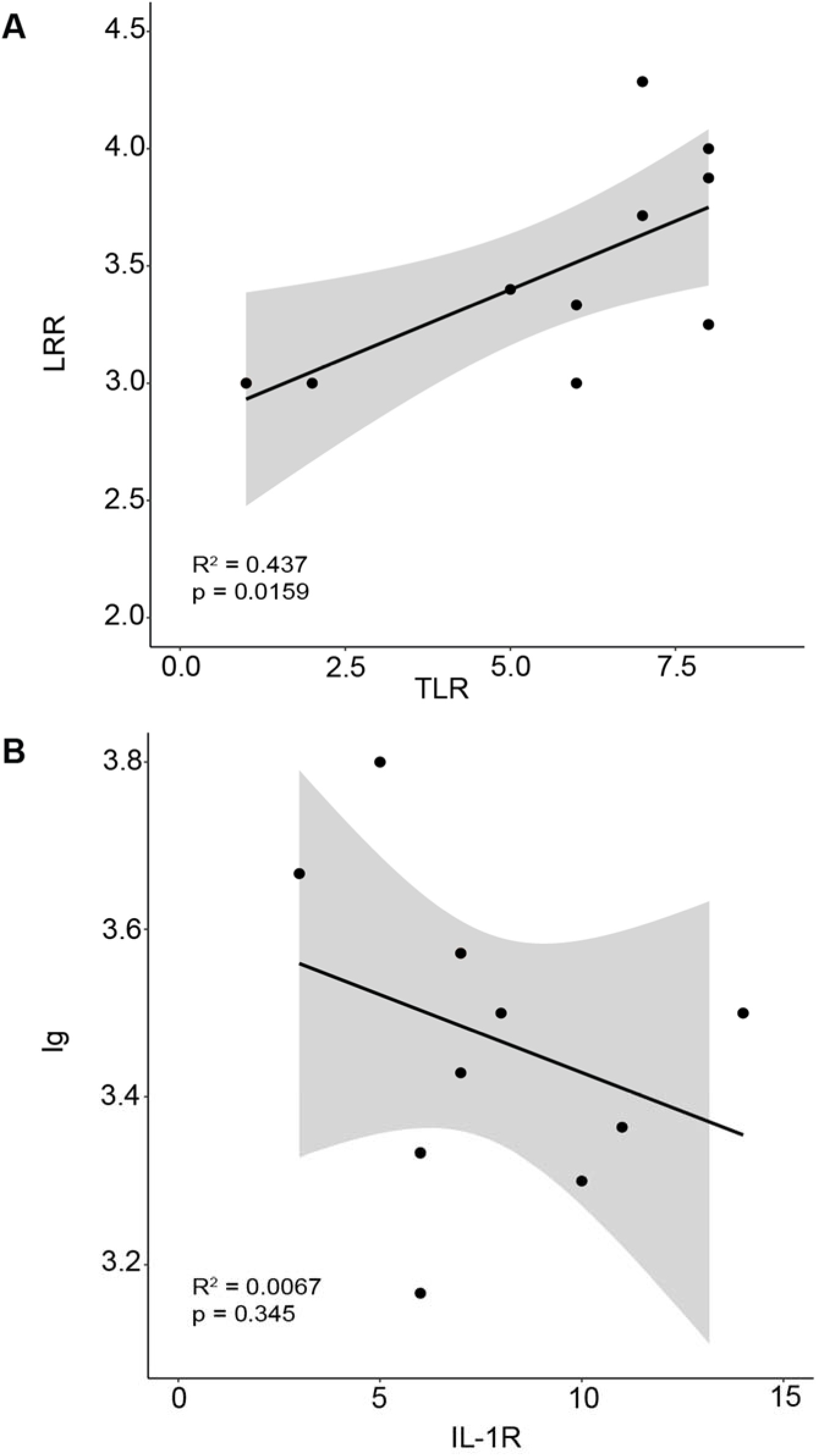
Comparison of domain copy number within genes and gene copies within genomes. **A** LRR domains vs TLR and **B** Ig vs IL-1R plots.

**Fig. S10.**
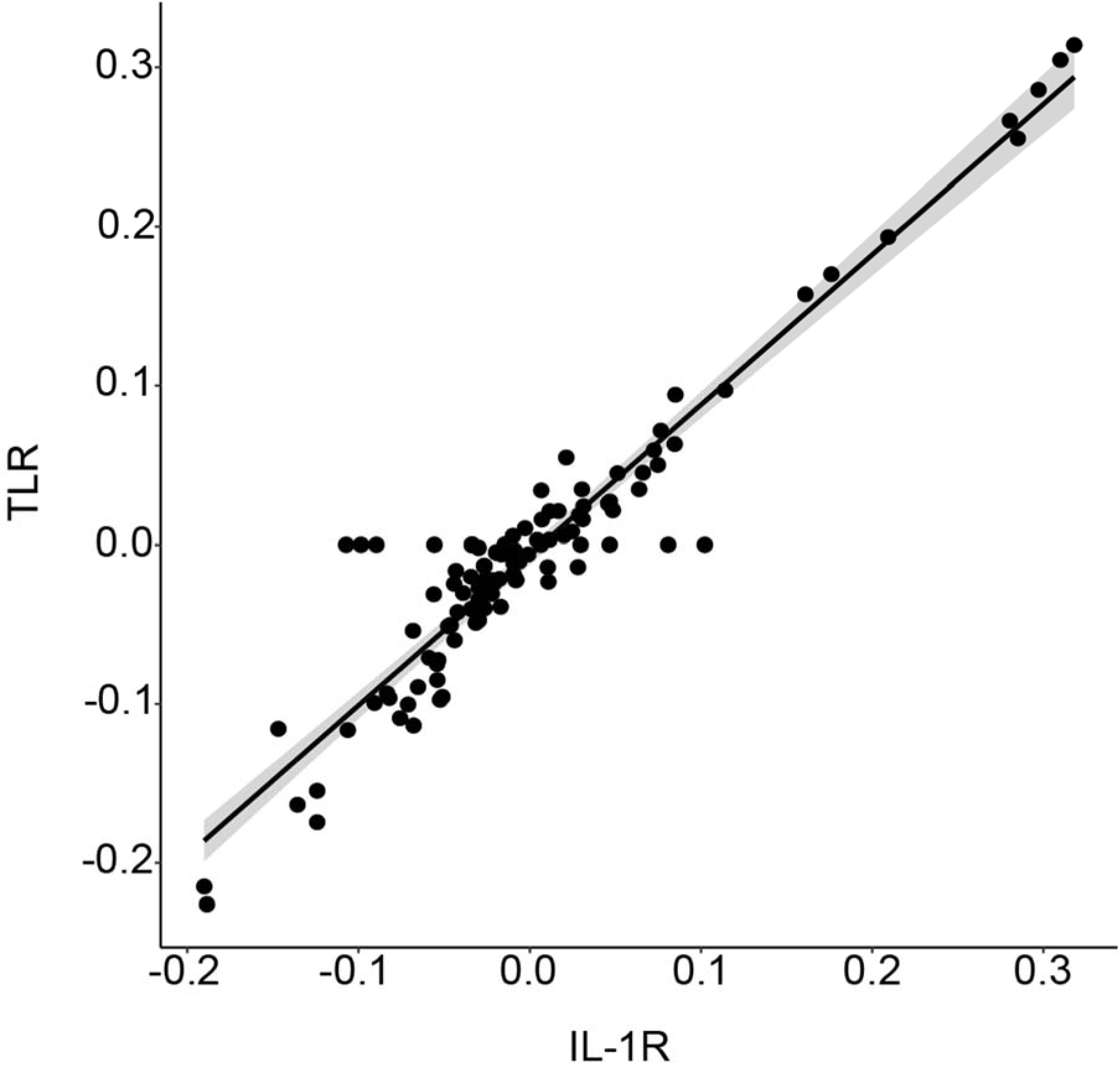
Correlated response of microbial families to IL-1R or TLR gene copy number over evolution. Scatterplot shows contrasts in the correlation between microbial family relative abundance and IL-1R gene copy number (x-axis) and TLR gene copy number (y-axis). Each point represents one microbial family. Microbial families whose relative abundance is equally correlated with IL-1R and TLR gene copy number therefore fall near the diagonal.

**Fig. S11.**
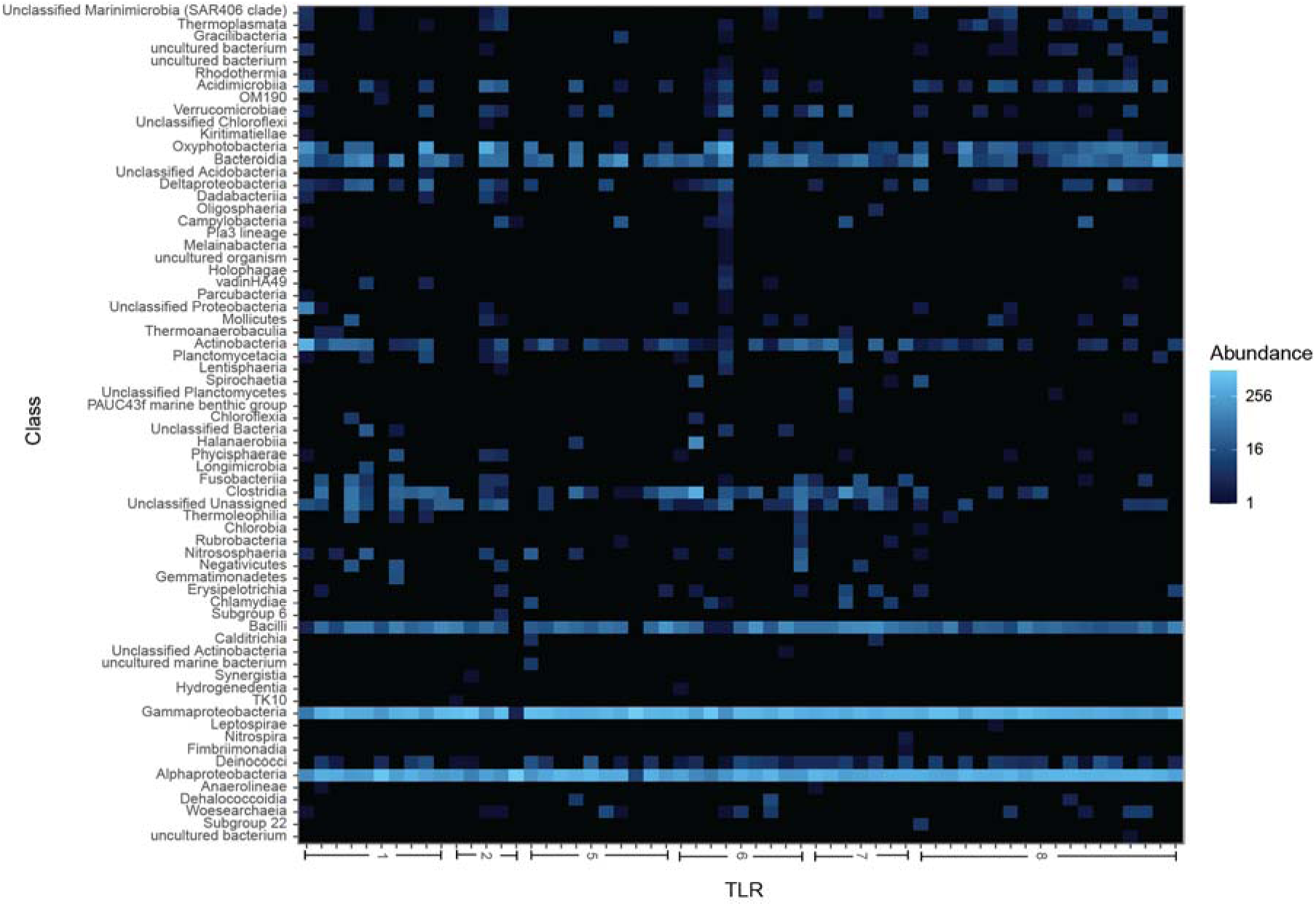
Heatmap of bacterial class abundance in mucus vs. TLR gene copy number. Heatmap colors depict the relative abundance of bacterial classes in coral mucus (y-axis), in samples sorted by TLR gene copy number (x-axis). Colors reflect abundance per 1000 16S rRNA gene amplicon reads.

**Fig. S12.**
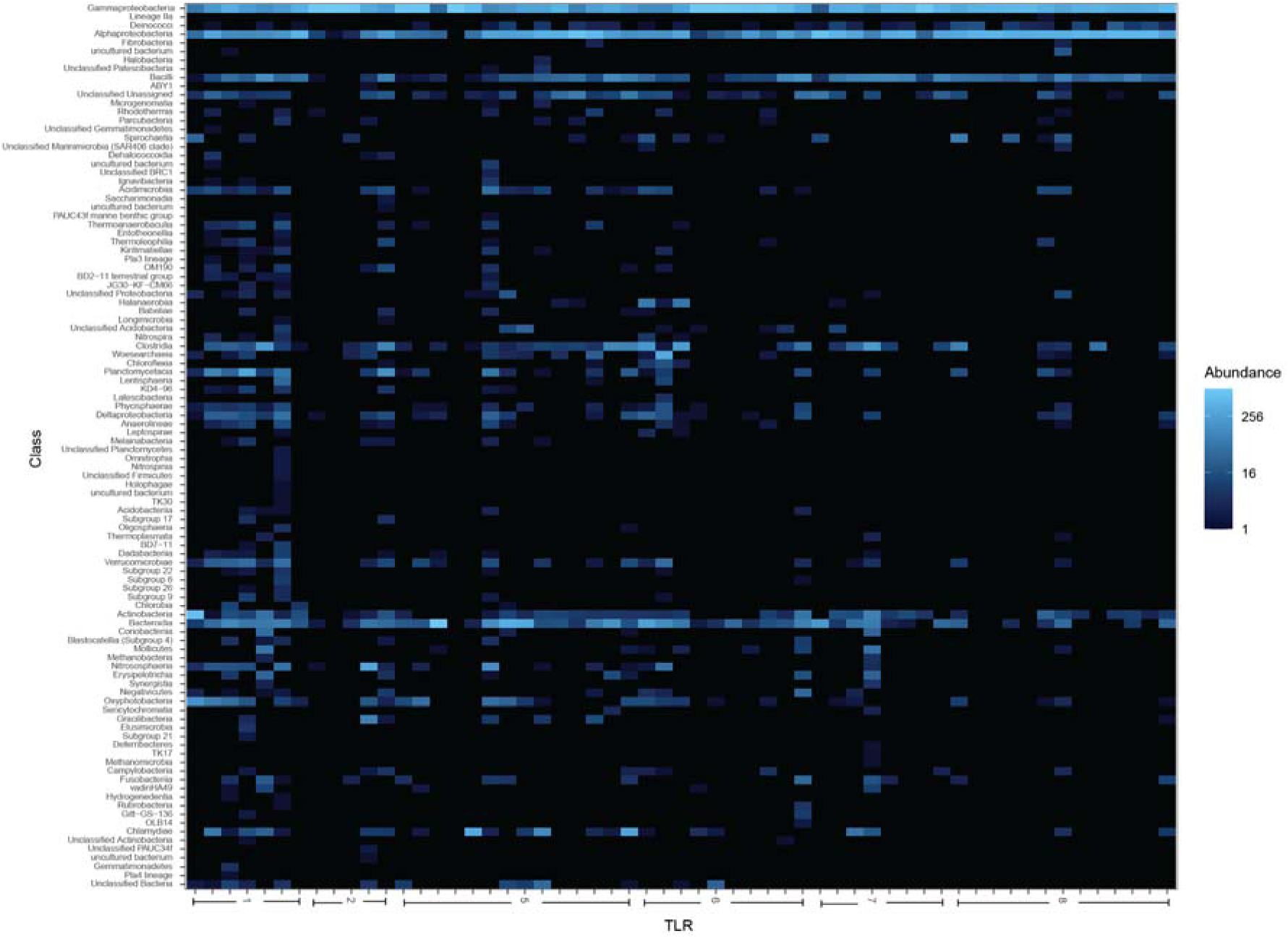
Heatmap of bacterial class relative abundance in tissue vs. TLR gene copy number. Heatmap colors depicts the relative abundance of bacterial classes in coral tissue (y-axis), in samples sorted by TLR gene copy number (x-axis). Colors reflect abundance per 1000 16S rRNA gene amplicon reads.

**Fig. S13.**
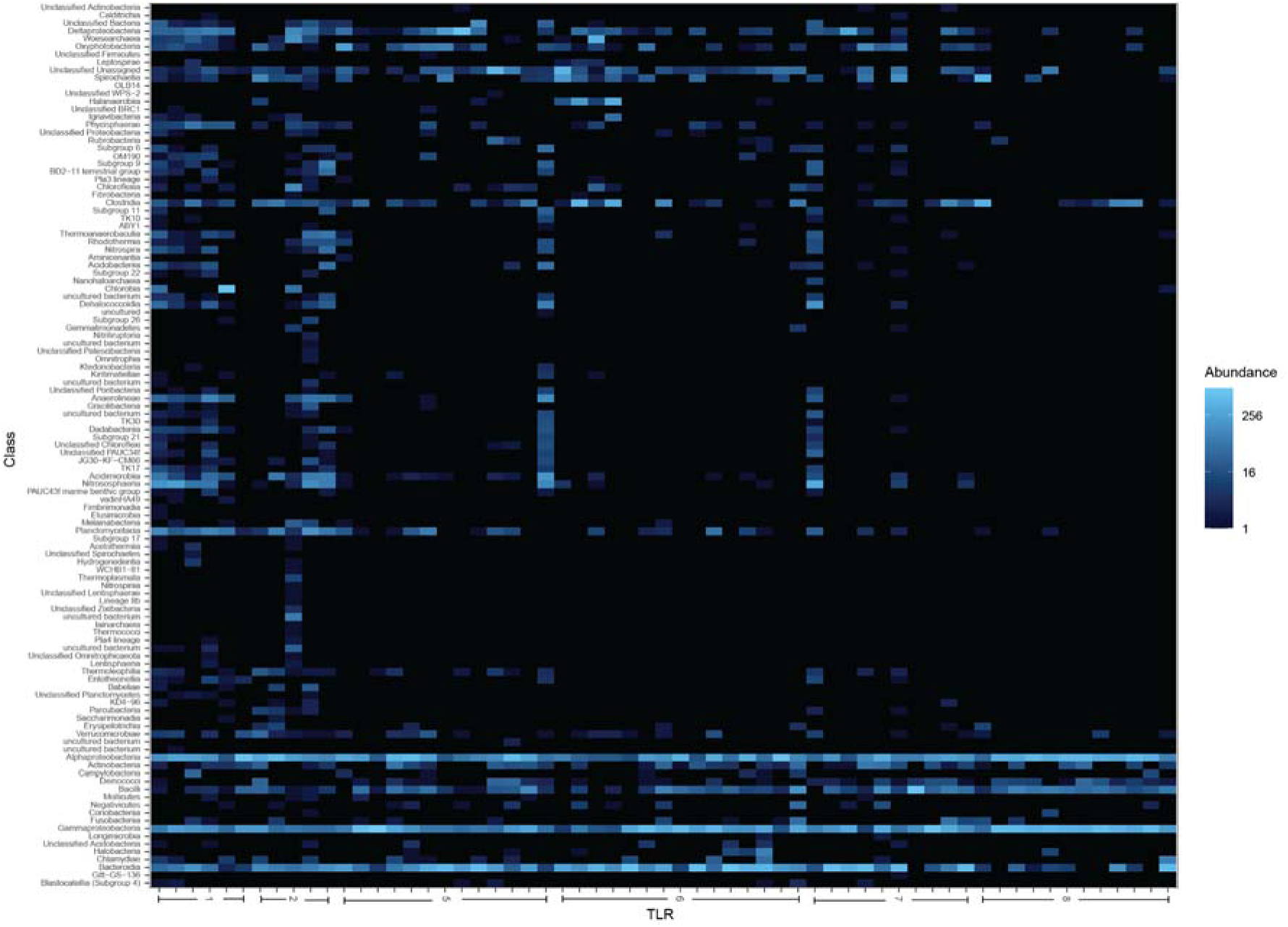
Heatmap of bacterial class relative abundance in skeleton vs. TLR gene copy number. Heatmap colors depict the relative abundance of bacterial classes in coral skeleton (y-axis), in samples sorted by TLR gene copy number (x-axis). Colors reflect abundance per 1000 16S rRNA gene amplicon reads.

**Fig. S14.**
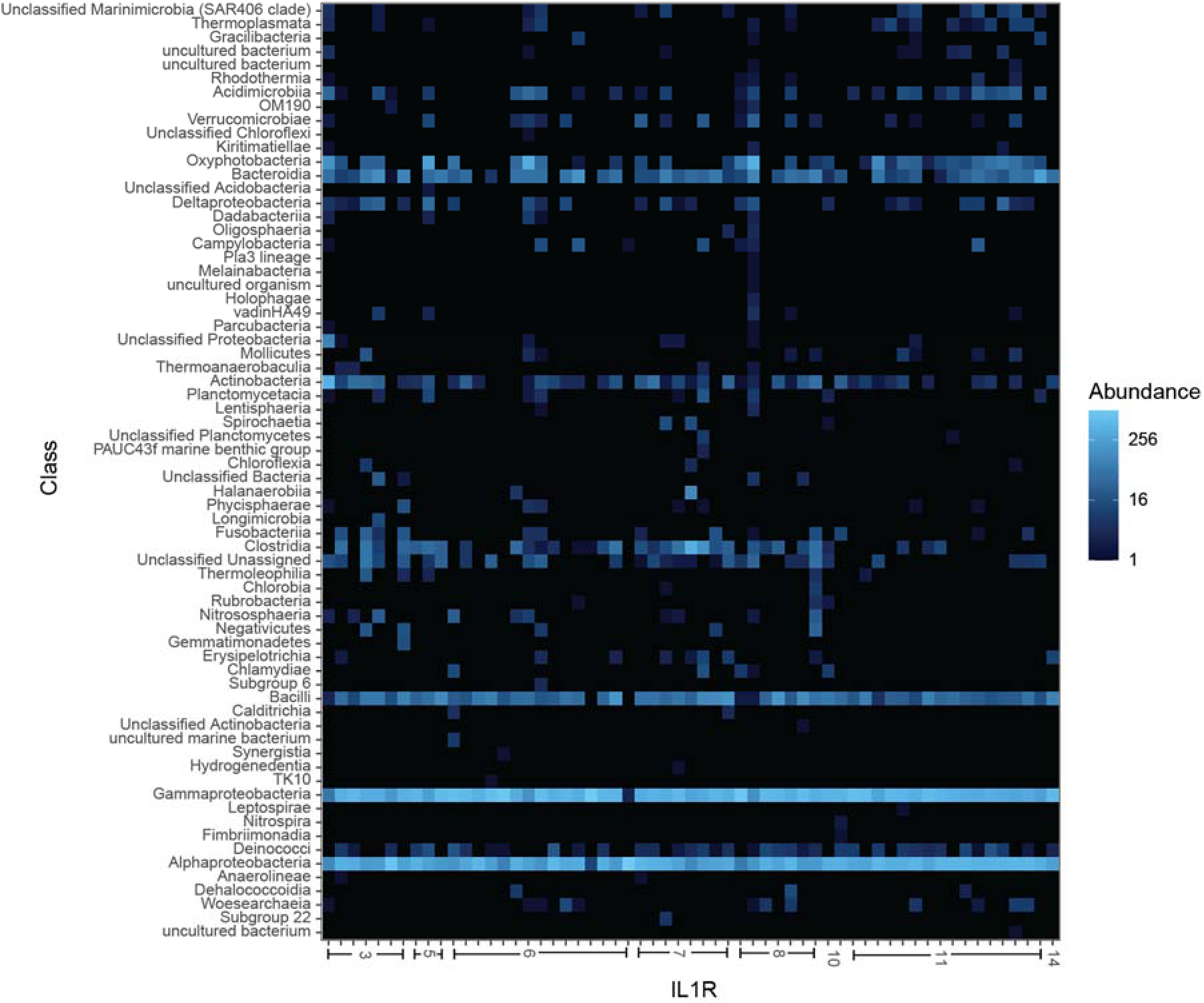
Heatmap of bacterial class abundance in mucus vs. IL-1R gene copy number. Heatmap colors depict the relative abundance of bacterial classes in coral mucus (y-axis), in samples sorted by IL-1R gene copy number (x-axis). Colors reflect abundance per 1000 16S rRNA gene amplicon reads.

**Fig. S15.**
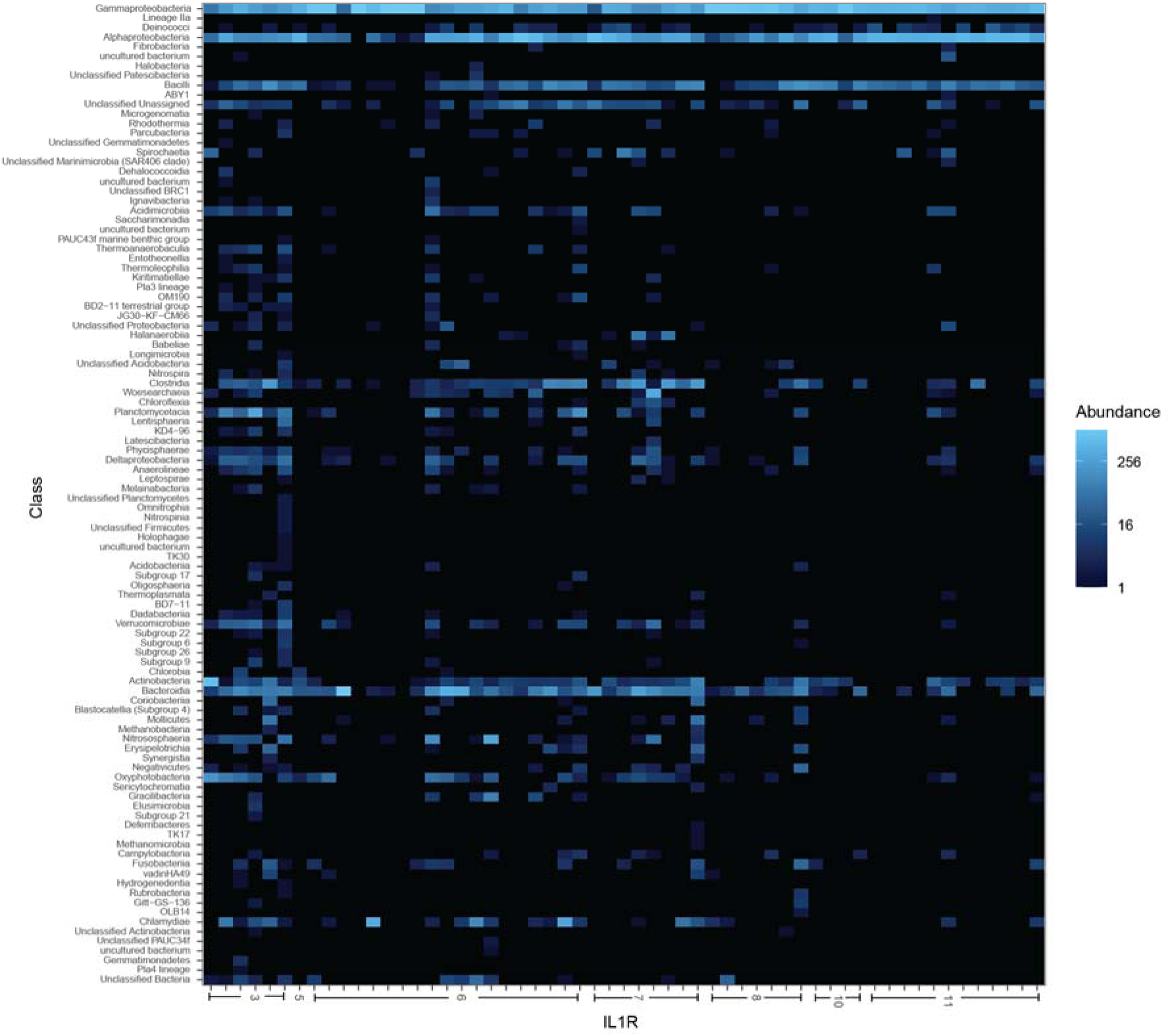
Heatmap of bacterial class abundance in tissue vs. IL-1R gene copy number. Heatmap colors depict the relative abundance of bacterial classes in coral tissue (y-axis), in samples sorted by IL-1R gene copy number (x-axis). Colors reflect abundance per 1000 16S rRNA gene amplicon reads.

**Fig. S16.**
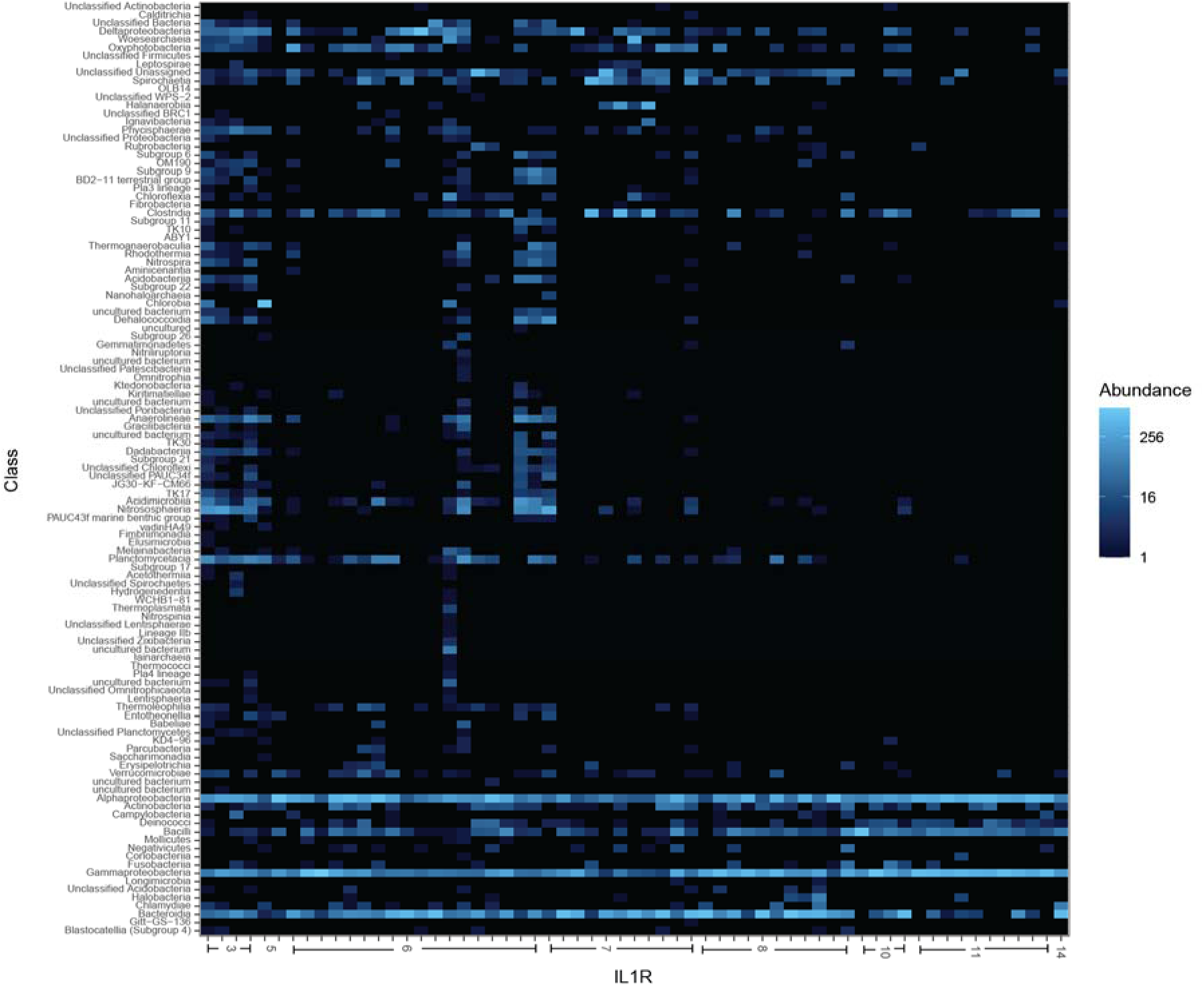
Heatmap of bacterial class abundance in skeleton vs. IL-1R gene copy number. Heatmap colors depict the relative abundance of bacterial classes in coral skeleton (y-axis), in samples sorted by IL-1R gene copy number (x-axis). Colors reflect abundance per 1000 16S rRNA gene amplicon reads.

## Supplementary Data Tables

**Supplementary Table 1.** Sample and Genomic metadata. (**A**) Genomes used in the analysis (B) Per sample metadata for the GCMP data (**C**) Mapping file for the GCMP data.

**Supplementary Table 2.** Annotations of TIR-domain containing gene families in coral genomes for (**A**) Total number of IL-1R and TLR genes, (**B**) Domain make up of IL-1R genes, (**C**) Domain make up of TLR genes.

**Supplementary Table 3.** Quality control information for the sequencing depth of microbiome samples in the dataset.

**Supplementary Table 4.** Distribution of TIR, LRR, and Ig domains across unique predicted protein isoforms in sequenced coral genomes.

**Supplementary Table 5.** Microbiome (**A**) richness and (**B**) evenness across samples in the analysis.

**Supplementary Table 6**. Phylogenetic comparison of IL-1R and TLR gene family copy number vs. microbiome alpha diversity in (**A**) All Samples (**B**) Mucus (**C**) Tissue, (**D**) Skeleton.

**Supplementary Table 7.** Trait table of the genomic and microbiome traits of coral species in the analysis.

**Supplementary Table 8.** Phylogenetic comparison of TIR LRR and Ig domains vs. microbiome alpha diversity in A) All Samples B) Mucus c) Tissue, d) Skeleton.

**Supplementary Table 9**. Phylogenetic comparison of IL-1R and TLR gene family copy number vs. PC axes from beta-diversity PCoA analysis in (**A**) All Samples (**B**) Mucus

(**C**) Tissue, (**D**) Skeleton.

**Supplementary Table 10.** ANCOMBC comparison of relative abundance of microbial taxa vs. IL-1R and TLR gene family copy number in (**A**) All Samples (**B**) Mucus (**C**) Tissue, (**D**) Skeleton.

**Supplementary Table 11.** Phylogenetic comparison of TIR LRR and Ig domains vs. microbiome PC axes from beta-diversity PCoA analysis in (**A**) All Samples (**B**) Mucus (**C**) Tissue, (**D**) Skeleton.

## References

1. Hoeksema, B. W. & Cairns, S. World List of Scleractinia. https://www.marinespecies.org/scleractinia.

2. Rohwer, F., Seguritan, V., Azam, F. & Knowlton, N. Diversity and distribution of coral-associated bacteria. Mar. Ecol. Prog. Ser. 243, 1–10 (2002).

3. Pollock, F. J. et al. Coral-associated bacteria demonstrate phylosymbiosis and cophylogeny. Nat. Commun. 9, 4921 (2018).

4. Ricci, F. et al. Host Traits and Phylogeny Contribute to Shaping Coral-Bacterial Symbioses. Msystems 7, e00044–22 (2022).

5. Hernandez-Agreda, A., Leggat, W., Bongaerts, P., Herrera, C. & Ainsworth, T. D. Rethinking the coral microbiome: simplicity exists within a diverse microbial biosphere. MBio 9, e00812–18 (2018).

6. Littman, R. A., Willis, B. L., Pfeffer, C. & Bourne, D. G. Diversities of coral-associated bacteria differ with location, but not species, for three acroporid corals on the Great Barrier Reef. FEMS Microbiol. Ecol. 68, 152–163 (2009).

7. Rodriguez-Lanetty, M., Granados-Cifuentes, C., Barberan, A., Bellantuono, A. J. & Bastidas, C. Ecological Inferences from a deep screening of the C omplex B acterial C onsortia associated with the coral, *Porites astre*oides. Mol. Ecol. 22, 4349–4362 (2013).

8. Pantos, O., Bongaerts, P., Dennis, P. G., Tyson, G. W. & Hoegh-Guldberg, O. Habitat-specific environmental conditions primarily control the microbiomes of the coral *Seriatopora hystrix*. ISME J. 9, 1916–1927 (2015).

9. Dunphy, C. M., Gouhier, T. C., Chu, N. D. & Vollmer, S. V. Structure and stability of the coral microbiome in space and time. Sci. Rep. 9, 1–13 (2019).

10. Wang, L. et al. Corals and their microbiomes are differentially affected by exposure to elevated nutrients and a natural thermal anomaly. Front. Mar. Sci. 5, 101 (2018).

11. Pootakham, W. et al. Heat-induced shift in coral microbiome reveals several members of the Rhodobacteraceae family as indicator species for thermal stress in Porites lutea. MicrobiologyOpen 8, e935 (2019).

12. Ricci, F. et al. Beneath the surface: community assembly and functions of the coral skeleton microbiome. Microbiome 7, 1–10 (2019).

13. Kusdianto, H. et al. Microbiomes of Healthy and Bleached Corals During a 2016 Thermal Bleaching Event in the Upper Gulf of Thailand. Front. Mar. Sci. 8, 643962 (2021).

14. Marchioro, G. M. et al. Microbiome dynamics in the tissue and mucus of acroporid corals differ in relation to host and environmental parameters. PeerJ 8, e9644 (2020).

15. Fifer, J. E. et al. Microbiome structuring within a coral colony and along a sedimentation gradient. Cell. Stress Response Physiol. Adapt. Corals Subj. Environ. Stress. Pollut. (2022).

16. Zaneveld, J. R. et al. Overfishing and nutrient pollution interact with temperature to disrupt coral reefs down to microbial scales. Nat. Commun. 7, 1–12 (2016).

17. Corinaldesi, C. et al. Multiple impacts of microplastics can threaten marine habitat-forming species. *Commun*. Biol. 4, 1–13 (2021).

18. Pratte, Z. A., Longo, G. O., Burns, A. S., Hay, M. E. & Stewart, F. J. Contact with turf algae alters the coral microbiome: contact versus systemic impacts. Coral Reefs 37, 1–13 (2018).

19. Briggs, A. A., Brown, A. L. & Osenberg, C. W. Local versus site-level effects of algae on coral microbial communities. R. Soc. Open Sci. 8, 210035 (2021).

20. Miller, N., Maneval, P., Manfrino, C., Frazer, T. K. & Meyer, J. L. Spatial distribution of microbial communities among colonies and genotypes in nursery-reared *Acropora cervicornis*. PeerJ 8, e9635 (2020).

21. Palacio-Castro, A. M., Rosales, S. M., Dennison, C. E. & Baker, A. C. Microbiome signatures in *Acropora cervicornis* are associated with genotypic resistance to elevated nutrients and heat stress. Coral Reefs 41, 1389–1403 (2022).

22. Rosales, S. M. et al. Microbiome differences in disease-resistant vs. susceptible Acropora corals subjected to disease challenge assays. Sci. Rep. 9, 18279 (2019).

23. Rosado, P. M. et al. Marine probiotics: increasing coral resistance to bleaching through microbiome manipulation. ISME J. 13, 921–936 (2019).

24. Ley, R. E. et al. Evolution of mammals and their gut microbes. science 320, 1647–1651 (2008).

25. Bodawatta, K. H. et al. Species-specific but not phylosymbiotic gut microbiomes of New Guinean passerine birds are shaped by diet and flight-associated gut modifications. Proc. R. Soc. B 288, 20210446 (2021).

26. Poole, A. Z. & Weis, V. M. TIR-domain-containing protein repertoire of nine anthozoan species reveals coral–specific expansions and uncharacterized proteins. Dev. Comp. Immunol. 46, 480–488 (2014).

27. Medzhitov, R. & Janeway, C. A. Innate immunity: the virtues of a nonclonal system of recognition. Cell 91, 295–298 (1997).

28. Thompson, L. R. et al. A communal catalogue reveals Earth’s multiscale microbial diversity. Nature 551, 457–463 (2017).

29. Parada, A. E., Needham, D. M. & Fuhrman, J. A. Every base matters: assessing small subunit rRNA primers for marine microbiomes with mock communities, time series and global field samples. Environ. Microbiol. 18, 1403–1414 (2016).

30. Apprill, A., McNally, S., Parsons, R. & Weber, L. Minor revision to V4 region SSU rRNA 806R gene primer greatly increases detection of SAR11 bacterioplankton. Aquat. Microb. Ecol. 75, 129–137 (2015).

31. Caporaso, J. G. EMP 16S Illumina Amplicon Protocol. (2018) doi:10.17504/protocols.io.nuudeww.

32. Gonzalez, A. et al. Qiita: rapid, web-enabled microbiome meta-analysis. Nat. Methods 15, 796–798 (2018).

33. Caporaso, J. G. et al. QIIME allows analysis of high-throughput community sequencing data. Nat. Methods 7, 335–336 (2010).

34. Amir, A. et al. Deblur Rapidly Resolves Single-Nucleotide Community Sequence Patterns. mSystems 2, e00191–16 (2017).

35. Bolyen, E. et al. QIIME 2: Reproducible, interactive, scalable, and extensible microbiome data science. https://peerj.com/preprints/27295 (2018) doi:10.7287/peerj.preprints.27295v2.

36. Rognes, T., Flouri, T., Nichols, B., Quince, C. & Mahé, F. VSEARCH: a versatile open source tool for metagenomics. PeerJ 4, e2584 (2016).

37. Quast, C. et al. The SILVA ribosomal RNA gene database project: improved data processing and web-based tools. Nucleic Acids Res. 41, D590–D596 (2013).

38. Bengtsson-Palme, J. et al. METAXA2: improved identification and taxonomic classification of small and large subunit rRNA in metagenomic data. Mol. Ecol. Resour. 15, 1403–1414 (2015).

39. Sonett, D., Brown, T., Bengtsson-Palme, J., Padilla-Gamiño, J. L. & Zaneveld, J. R. The Organelle in the Room: Under-annotated Mitochondrial Reads Bias Coral Microbiome Analysis. http://biorxiv.org/lookup/doi/10.1101/2021.02.23.431501 (2021) doi:10.1101/2021.02.23.431501.

40. Janssen, S., et al. Phylogenetic Placement of Exact Amplicon Sequences Improves Associations with Clinical Information. mSystems 3, (2018).

41. Lin, H. & Peddada, S. D. Analysis of compositions of microbiomes with bias correction. Nat. Commun. 11, 1–11 (2020).

42. Emery, M. A., Dimos, B. A. & Mydlarz, L. D. Cnidarian pattern recognition receptor repertoires reflect both phylogeny and life history traits. Front. Immunol. 12, 689463 (2021).

43. Mazel, F. et al. Is host filtering the main driver of phylosymbiosis across the tree of life? Msystems 3, e00097–18 (2018).

44. Brooks, A. W., Kohl, K. D., Brucker, R. M., van Opstal, E. J. & Bordenstein, S. R. Phylosymbiosis: Relationships and Functional Effects of Microbial Communities across Host Evolutionary History. PLOS Biol. 14, e2000225 (2016).

45. O’Brien, P. A. et al. Diverse coral reef invertebrates exhibit patterns of phylosymbiosis. ISME J. 14, 2211–2222 (2020).

46. Delsuc, F. et al. Convergence of gut microbiomes in myrmecophagous mammals. Mol. Ecol. 23, 1301–1317 (2014).

47. Song, S. J. et al. Is there convergence of gut microbes in blood-feeding vertebrates? Philos. Trans. R. Soc. B Biol. Sci. 374, 20180249 (2019).

48. Song, S. J. et al. Comparative Analyses of Vertebrate Gut Microbiomes Reveal Convergence between Birds and Bats. mBio 11, e02901–19 (2020).

49. Palmer, C. V. et al. Patterns of coral ecological immunology: variation in the responses of Caribbean corals to elevated temperature and a pathogen elicitor. J. Exp. Biol. 214, 4240–4249 (2011).

50. Palmer, C., Bythell, J. & Willis, B. Enzyme activity demonstrates multiple pathways of innate immunity in Indo-Pacific anthozoans. Proc. R. Soc. B Biol. Sci. 279, 3879–3887 (2012).

51. Brown, T., Sonett, D., Zaneveld, J. R. & Padilla-Gamiño, J. L. Characterization of the microbiome and immune response in corals with chronic Montipora white syndrome. Mol. Ecol. 30, 2591–2606 (2021).

52. Thaiss, C. A., Levy, M., Suez, J. & Elinav, E. The interplay between the innate immune system and the microbiota. Curr. Opin. Immunol. 26, 41–48 (2014).

53. Fonseca, J. P., Lakshmanan, V., Boschiero, C. & Mysore, K. S. The Pattern Recognition Receptor FLS2 Can Shape the Arabidopsis Rhizosphere Microbiome β-Diversity but Not EFR1 and CERK1. Plants 11, 1323 (2022).

54. Axworthy, J. B. et al. Shotgun Proteomics Identifies Active Metabolic Pathways in Bleached Coral Tissue and Intraskeletal Compartments. Front. Mar. Sci. (2022).

55. Levy, S. & Mass, T. The Skeleton and Biomineralization Mechanism as Part of the Innate Immune System of Stony Corals. Front. Immunol. 13, (2022).

56. Vega Thurber, R., et al. Deciphering coral disease dynamics: integrating host, microbiome, and the changing environment. Front. Ecol. Evol. 8, 575927 (2020).

57. Elton, C. S. The Ecology of Invasions by Animals and Plants. (Springer Nature, 2020).

58. Marraffini, M. L. & Geller, J. B. Species richness and interacting factors control invasibility of a marine community. Proc. R. Soc. B Biol. Sci. 282, 20150439 (2015).

59. Sommer, F. et al. Microbiomarkers in inflammatory bowel diseases: caveats come with caviar. Gut 66, 1734–1738 (2017).

60. Sommer, F., Anderson, J. M., Bharti, R., Raes, J. & Rosenstiel, P. The resilience of the intestinal microbiota influences health and disease. Nat. Rev. Microbiol. 15, 630– 638 (2017).

61. Palmer, C. V. Immunity and the coral crisis. Commun. Biol. 1, 91 (2018).

62. Feng, G., Sun, W., Zhang, F., Karthik, L. & Li, Z. Inhabitancy of active Nitrosopumilus-like ammonia-oxidizing archaea and Nitrospira nitrite-oxidizing bacteria in the sponge Theonella swinhoei. Sci. Rep. 6, 1–11 (2016).

63. Kim, S.-H., Kim, J.-H., Park, M.-A., Hwang, S. D. & Kang, J.-C. The toxic effects of ammonia exposure on antioxidant and immune responses in Rockfish, Sebastes schlegelii during thermal stress. Environ. Toxicol. Pharmacol. 40, 954–959 (2015).

64. Kvennefors, E., Sampayo, E., Ridgway, T., Barnes, A. & Hoegh-Guldberg, O. Bacterial Communities of Two Ubiquitous Great Barrier Reef Corals. (2010).

65. Aronson, R. B. & Precht, W. F. White-band disease and the changing face of Caribbean coral reefs. in The ecology and etiology of newly emerging marine diseases 25–38 (Springer, 2001).

66. Casas, V. et al. Widespread association of a Rickettsiales-like bacterium with reef-building corals. Environ. Microbiol. 6, 1137–1148 (2004).

67. Gignoux-Wolfsohn, S., Marks, C. J. & Vollmer, S. V. White Band Disease transmission in the threatened coral, *Acropora cervicornis*. Sci. Rep. 2, 1–3 (2012).

68. Miller, M. W., Lohr, K. E., Cameron, C. M., Williams, D. E. & Peters, E. C. Disease dynamics and potential mitigation among restored and wild staghorn coral, *Acropora cervicornis*. PeerJ 2, e541 (2014).

69. Wan, S. J. et al. IL-1R and MyD88 Contribute to the Absence of a Bacterial Microbiome on the Healthy Murine Cornea. Front. Microbiol. 9, 1117 (2018).

70. Gerardo, N. M., Hoang, K. L. & Stoy, K. S. Evolution of animal immunity in the light of beneficial symbioses. Philos. Trans. R. Soc. Lond. B. Biol. Sci. 375, 20190601 (2020).

71. McFall-Ngai, M. Care for the community. Nature 445, 153–153 (2007).

72. Moreira, L. A. et al. A Wolbachia symbiont in Aedes aegypti limits infection with dengue, Chikungunya, and Plasmodium. Cell 139, 1268–1278 (2009).

73. Lee, K. H. & Ruby, E. G. Effect of the Squid Host on the Abundance and Distribution of Symbiotic Vibrio fischeri in Nature. Appl. Environ. Microbiol. 60, 1565– 1571 (1994).

74. Visick, K. L., Foster, J., Doino, J., McFall-Ngai, M. & Ruby, E. G. *Vibrio fischeri* lux genes play an important role in colonization and development of the host light organ. J. Bacteriol. 182, 4578–4586 (2000).

75. Broderick, N. A. Friend, foe or food? Recognition and the role of antimicrobial peptides in gut immunity and Drosophila–microbe interactions. Philos. Trans. R. Soc. B Biol. Sci. 371, (2016).

76. Kwong, W. K., Engel, P., Koch, H. & Moran, N. A. Genomics and host specialization of honey bee and bumble bee gut symbionts. Proc. Natl. Acad. Sci. 111, 11509–11514 (2014).

77. Hillman, K. & Goodrich-Blair, H. Are you my symbiont? Microbial polymorphic toxins and antimicrobial compounds as honest signals of beneficial symbiotic defensive traits. Curr. Opin. Microbiol. 31, 184–190 (2016).

78. Fuess, L. E. et al. Immune Gene Expression Covaries with Gut Microbiome Composition in Stickleback. mBio 12, e00145–21 (2021).

79. Davies, C. S. et al. Immunogenetic variation shapes the gut microbiome in a natural vertebrate population. Microbiome 10, 41 (2022).

80. Evans, J. D. et al. Immune pathways and defence mechanisms in honey bees Apis mellifera. INSECT Mol. Biol. 15, 645–656 (2006).

81. Nyholm, S. V. & Graf, J. Knowing your friends: invertebrate innate immunity fosters beneficial bacterial symbioses. Nat. Rev. Microbiol. 10, 815–827 (2012).

